# Targeting the Schwann Cell EP2/cAMP Nanodomain to Block Pain but not Inflammation

**DOI:** 10.1101/2024.09.10.612200

**Authors:** Romina Nassini, Lorenzo Landini, Matilde Marini, Martina Chieca, Daniel Souza Monteiro de Araújo, Marco Montini, Pasquale Pensieri, Vittorio Donato Abruzzese, Gaetano De Siena, Jin Zhang, Elisa Bellantoni, Vincenzo De Giorgi, Antonia Romitelli, Giulia Brancolini, Raquel Tonello, Chloe J. Peach, Alessandra Mastricci, Irene Scuffi, Martina Tesi, Dane D. Jensen, Brian L. Schmidt, Nigel W. Bunnett, Francesco De Logu, Pierangelo Geppetti

## Abstract

Analgesia by non-steroidal anti-inflammatory drugs (NSAIDs) is ascribed to inhibition of prostaglandin (PG) biosynthesis and ensuing inflammation. However, NSAIDs have life-threatening side effects, and inhibition of inflammation delays pain resolution. Decoupling the mechanisms underlying PG-evoked pain *vs.* protective inflammation would facilitate pain treatment. Herein, we reveal that selective silencing of the PGE_2_ EP2 receptor in Schwann cells *via* an adeno-associated viral vector abrogates the indomethacin-sensitive component of pain-like responses in mice elicited by inflammatory stimuli without affecting inflammation. In human Schwann cells and in mice, EP2 activation and optogenetic stimulation of adenylyl cyclase evokes a plasma membrane-compartmentalized cyclic adenosine monophosphate (cAMP) signal that, *via* A-kinase anchor protein-associated protein kinase A, sustains inflammatory pain-like responses, but does not delay their resolution. Thus, an unforeseen and druggable EP2 receptor in Schwann cells, *via* specific cAMP nanodomains, encodes PG-mediated persistent inflammatory pain but not protective inflammation.

## Introduction

Inflammation protects against xenobiotics and mechanical, thermal and chemical injury^1^. Progression and resolution of inflammation are tightly regulated by pro- and anti-inflammatory factors^1^, including cyclooxygenase (COX)-byproducts, prostaglandins (PGs)^2^. Subcutaneous injection in rodents of PGE_2_, the major proalgesic and proinflammatory PG isoform^3^, evokes a transient non-evoked nociceptive response and a delayed and sustained mechanical hypersensitivity, attributed to direct stimulation of terminals of dorsal root ganglia (DRG) nociceptors^4,5^, *via* cyclic adenosine monophosphate (cAMP) formation and protein kinase A (PKA) activation^6^. A-kinase anchor protein (AKAP79/150)-dependent PKA also contributes to PGE_2_-induced nociceptor hypersensitivity^7^. Of the four PGE_2_ receptor subtypes (EP1-4), EP4 has been implicated in PGE_2_-induced excitation and sensitization of nociceptors^8^. However, conflicting data on the contribution of EP4 to carrageenan- and CFA-evoked mechanical hypersensitivity^9,10^, the failure of EP4-selective antagonist to provide benefit in migraine pain^11^, and absence of successful clinical trials with EP4 antagonists in inflammatory pain cast doubts on the PGE_2_ receptor subtype and its cellular and molecular pathways implicated in inflammatory pain.

SCs make a major contribution to the maintenance of mechanical allodynia in murine models of neuropathic^12^ and cancer^13^ pain. The promigraine neuropeptide, calcitonin gene-related peptide (CGRP)^14^ released from periorbital trigeminal nerve terminals elicits local mechanical allodynia by activating the calcitonin-like receptor and receptor activity-modifying protein 1 (CLR/RAMP1) on adjacent SCs, where sustained production cAMP in endosomes elicits persistent pain^15^. Disruption of low-density lipoprotein-receptor related protein-1 (LRP1) in SCs enhanced mechanical allodynia caused by nerve injury in mice^16^. Optogenetic stimulation of mouse cutaneous mouse SCs elicits mechanical hypersensitivity^17^. Herein, we report that AAV cell-selective silencing of SC EP2 abrogates PG-dependent mechanical allodynia and grimace behavior without affecting inflammation. EP2 in human and murine SCs activates a proalgesic signaling pathway encompassing plasma membrane-compartmentalized cAMP/PKA that, *via* AKAP79/150 anchoring to specific subcellular nanodomains, signals mechanical allodynia elicited by carrageen and CFA. This pathway is mechanistically independent from the PG-mediated inflammatory responses produced by these stimuli. In contrast with NSAIDs, SC EP2 antagonism or silencing reduces inflammatory pain without delaying its resolution.

## Results

### EP2 mediates sustained PGE_2_-evoked mechanical allodynia

The identity of the PGE_2_ receptor subtype(s) involved in inflammatory pain remains elusive. Studies with global receptor knockout mice or systemically administered receptor antagonists have provided confounding results because of opposing activities of central *vs.* peripheral receptors^9^ and incomplete antagonist selectivity^18^. To find the peripheral EP subtype(s) implicated in PG-dependent inflammatory pain, agents were administered by local intraplantar (i.pl.) injection into the mouse hindpaw (unless otherwise specified), and to identify the receptor cellular localization EP subtypes were selectively silenced in SCs or DRG neurons. In C57BL/6J (B6) mice injected with PGE_2_, we assessed the two temporally distinct responses reported in previous studies^3–5^: the early and transient (∼15 min) non-evoked nociception, consisting of lifting and licking, but not shaking and biting (Figure 1a and Supplementary Fig.1a), and a delayed (∼30 min) and sustained (∼4 hours) hindpaw mechanical allodynia (allodynia) (Figure 1b), which were observed solely in the injected paw (Supplementary Fig.1b), indicating a locally confined action of PGE_2_. As no difference was found between male and female mice (Supplementary Fig.1c,d), to minimize the number of animals used, only key experiments were replicated in both sexes. The selective EP4 agonist, L-902,688 (L-902)^19^, elicited an early and transient non-evoked nociception and a moderate delayed allodynia (Figure 1c-e). In contrast, the selective EP2 agonist, butaprost^20^, elicited a robust allodynia, without detectable non-evoked nociception (Figure 1f,g). The EP2 antagonist PF-04418948 (PF)^21^ reduced allodynia evoked by PGE_2_ and butaprost without affecting non-evoked nociception to PGE_2_ and L-902 (Figure 1h,i; Supplementary Fig.1e,f). The EP4 antagonist BGC20-1531 (BGC)^22^, produced marginal or no reduction of PGE_2_- or butaprost-evoked allodynia, respectively, whereas it dose-dependently abolished non-evoked nociception to PGE_2_ and L-902 (Fig.1h,i; Supplementary Fig.1g-k). The role of EP4 in PGE_2_-induced allodynia^23^, studied using the putative selective antagonist, ER819762 (ER81),^24^ was not confirmed here. Although ER81 reduced PGE_2_- and L-902-induced non-evoked nociception and allodynia, unlike BGC, higher doses of ER81 unselectively attenuated allodynia by the EP2 selective agonist butaprost (Supplementary Fig.1l-p). Therefore, previous confounding results are attributable to incomplete selectivity of early EP antagonists. EP1 has been implicated in PGE_2_-induced heat hyperalgesia^25^. However, neither EP1 (SC-51322) nor EP3 (L-798,106) antagonists affected non-evoked nociception or allodynia to PGE_2_ (Supplementary Fig.1q-t). Similarly, prostacyclin receptor (IP) antagonism (Ro-1138452) neither reduced non-evoked nociception nor allodynia to PGE_2_ (Supplementary Fig.1u,v). Thus, EP4 alone mediates the early and transient nociception, whereas EP2 primarily mediates the delayed and sustained allodynia induced by PGE_2_.

**Figure 1.**
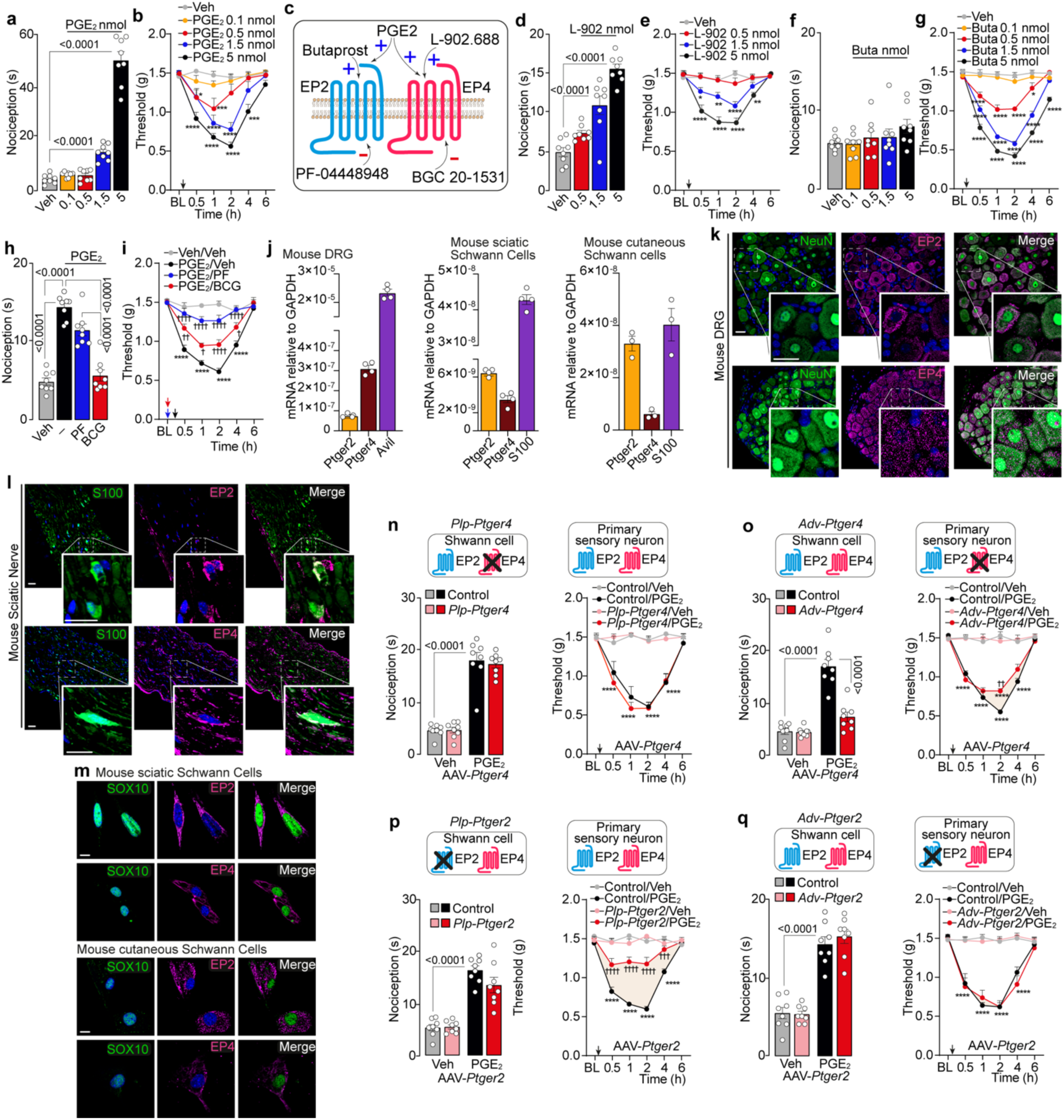
EP2 receptor activation in Schwann cells mediates PGE_2_-evoked sustained hindpaw mechanical allodynia (allodynia). **a, d, f,** Dose-dependent acute nociception and **b, e, g,** dose- and time-dependent allodynia after intraplantar (i.pl.) injection of PGE_2_, L-902,688 (L-902), butaprost (Buta) or vehicle (Veh) in C57BL/6J mice (B6). **c,** Schematic representation of agonists/antagonists targeting EP2 and EP4 receptors. **h,** Acute nociception and i, allodynia after i.pl. PGE_2_ (1.5 nmol) or Veh in B6 pretreated with PF-04448948 (PF, 5nmol), BGC 20-1531 (BGC, 5 nmol) or Veh (n=8 mice per group). **j,** RT-qPCR for *Ptger2, Ptger4 Avil* and *S100* mRNA in mouse dorsal root ganglia (DRG) sciatic and cutaneous Schwann cells (SCs) (DRG and sciatic SCs n=4, cutaneous n=3 independent experiments). **k, l,** Representative images of EP2, EP4, NeuN and S100B expression in mouse DRG and sciatic nerve tissue (scale bar: 20 μm) (n=3 subjects). **m,** Representative images of EP2, EP4 and SOX10 expression in mouse sciatic and cutaneous SCs (n=3 independent experiments). **n-q,** Acute nociception (left panel) and allodynia (right panel) after i.pl. PGE_2_ or Veh in *Plp-Cre, Adv-Cre* or Control mice infected with AAV for selective silencing of EP4 *(-Ptger4*) (*P1p-Ptger4* or *Adv-Ptger4*) (**n,o**) or EP2 *(-Ptger2*) (*P1p-Ptger2* or *Adv-Ptger2*) (**p,q**) (n=8 mice per group). Data are mean ± s.e.m. **a,d, f, h, n, o, p, q** 1-way or **b, e, g, i, n, o, p**, q 2-way ANOVA, Bonferroni correction. *P<0.05, **P<0.01, ***P<0.001, ****P<0.0001 vs. Veh ^†^P<0.05, ^††^P<0.01, ^†††^P<0.001, ^††††^P<0.0001 vs. PGE_2_/Veh, Control/ PGE_2_.

### SC EP2 mediates sustained PGE_2_-evoked mechanical allodynia

The essential contribution of SCs to pain-like responses in preclinical models of neuropathic^12^, cancer^13^ and migraine^15^ pain was confirmed by optogenetic evidence ^17^. Whereas both direct and indirect data support the presence and function of EP2 and EP4 in DRG neurons^26^, far less is known about expression and function of these receptors in peripheral glial cells^27,28^. We detected EP2 and EP4 mRNA and protein in human and mouse DRG neurons and sciatic nerve SCs and in mouse cutaneous SCs (Supplementary Fig.2a-e; Figure 1j-m). To identify the cell type that mediates PGE_2_-induced non-evoked nociception and allodynia, we used *Plp^Cre^* or *Adv^Cre^* drivers to express short hairpin RNA (shRNA) in SCs of *Plp^Cre^* mice, or in DRG neurons of *Adv^Cre^*mice to selectively down- regulate EP2 or EP4 expression. Viruses were packaged with the AAV2/8 or AAV2/9n serotypes for efficient infection of SCs or DRG neurons, respectively, and in both cases with a loxP flanked shRNA of *Ptger2* (EP2) or *Ptger4* (EP4). Downregulation of EP2 or EP4 transcripts in sciatic nerve and DRG homogenates, and of EP2 or EP4 protein immunoreactivity in S100B (SCs) and NeuN (DRG neurons) positive cells, confirmed the efficiency and selectivity of shRNA-mediated silencing after intrasciatic (i.sc.) AAV2/8 injection in *Plp^Cre^* mice or intrathecal (i.th.) AAV2/9n injection in *Adv^Cre^* mice (Supplementary Fig.3a,b and Supplementary Fig.4a,b). No change in protein expression in non-targeted cells indicated selectivity (Supplementary Fig.3c). Expression of a floxed green fluorescence protein (GFP) reporter gene (control marker) also supported cell-selective targeting in *Plp^Cre^*or *Adv^Cre^* mice (Supplementary Fig.3d-f and Supplementary Fig.4c,d). Absence of inflammatory responses in AAV-shRNA-treated *Plp^Cre^* and control mice confirmed that AAV-shRNA does not *per se* cause inflammation (Supplementary Fig.3g). A scrambled shRNA (*scr-shRNA)* was used as control. In *Plp-scr-shRNA* mice, PGE_2_ evoked similar non-evoked nociception and allodynia as in control mice (Supplementary Fig.5a). To test efficiency of *scr-shRNA* expression, EP2 and EP4 and GFP protein were evaluated in S100B and NeuN positive cells (Supplementary Fig.5b-d). To exclude shRNA-dependent off-target effects, *Plp^Cre^* mice were infected with a shRNA directed to a different *Ptger2* region *(Plp-Ptger2-bis* mice). In *Plp-Ptger2-bis* mice, PGE_2_-dependent allodynia, but not non-evoked nociception, was reduced (Supplementary Fig.5e).

PGE_2_ evoked similar non-evoked nociception and allodynia in control and SC EP4 silenced (*Plp-Ptger4*) mice (Figure 1n). EP4 silencing in DRG neurons (*Adv-Ptger4* mice) inhibited markedly non-evoked nociception and only slightly allodynia in response to PGE_2_ (Figure 1o). In contrast, SC EP2 silencing (*Plp-Ptger2*) strongly inhibited PGE_2_-induced allodynia but not non-evoked nociception (Fig.1p). Non-evoked nociception and allodynia were not reduced in mice with EP2 silenced in DRG neurons (*Adv-Ptger2* mice) (Figure 1q). Non-evoked nociception and moderate allodynia elicited by L-902 (EP4 agonist) were attenuated in *Adv-Ptger4* mice (Supplementary Fig.5f,g), whereas the marked butaprost (EP2 agonist)-induced allodynia was attenuated in *Plp-Ptger2* mice (Supplementary Fig.5h). Non-evoked nociception and/or allodynia to L-902 or butaprost were not affected in any other silenced mouse strain (Supplementary Fig.5i-s). Thus, genetic silencing indicates that neuronal EP4 mediates non-evoked nociception whereas EP2 in SCs largely mediates mechanical allodynia induced by PGE_2_.

### SC EP2 mediates sustained carrageenan- and CFA-evoked mechanical allodynia and grimace behavior

To investigate the contribution of neuronal or glial EP4 and EP2 in pain-like responses elicited by endogenous PGs, we administered arachidonic acid (AA) and phospholipase A_2_ activating protein (PLAA), which are known to release eicosanoids^29^. AA and PLAA induced allodynia (∼4-6 h), without producing any measurable non-evoked nociceptive behavior (Figure 2a-d). AA and PLAA injection in B6 mouse hindpaw produced a slowly-developing increase in PGE_2_ hind paw levels that was abated by the COX inhibitor, indomethacin (Figure 2e,f). The PGE2 peak level was ∼50% of that produced by exogenous PGE_2_ injection (Figure 2e,f). The differential response to exogenous *vs.* endogenous PGE_2_ suggests that the non-evoked nociception by exogenous PGs is a pharmacological response that is not replicated by endogenously released PGs. SC EP2 silencing (*Plp-Ptger2* mice) (Figure 2g,h) or EP2 antagonism attenuated allodynia to AA or PLAA (Supplementary Fig.6a,b). Indomethacin treatment did not further enhance the attenuation produced by SC EP2 silencing (Figure 2g,h). All other interventions, including EP4 antagonism, SC and DRG neuron EP4 silencing and DRG EP2 silencing were ineffective (Supplementary Fig.6c-j), indicating a key role of the SC EP2 in sustaining allodynia by endogenous PGs.

**Figure 2.**
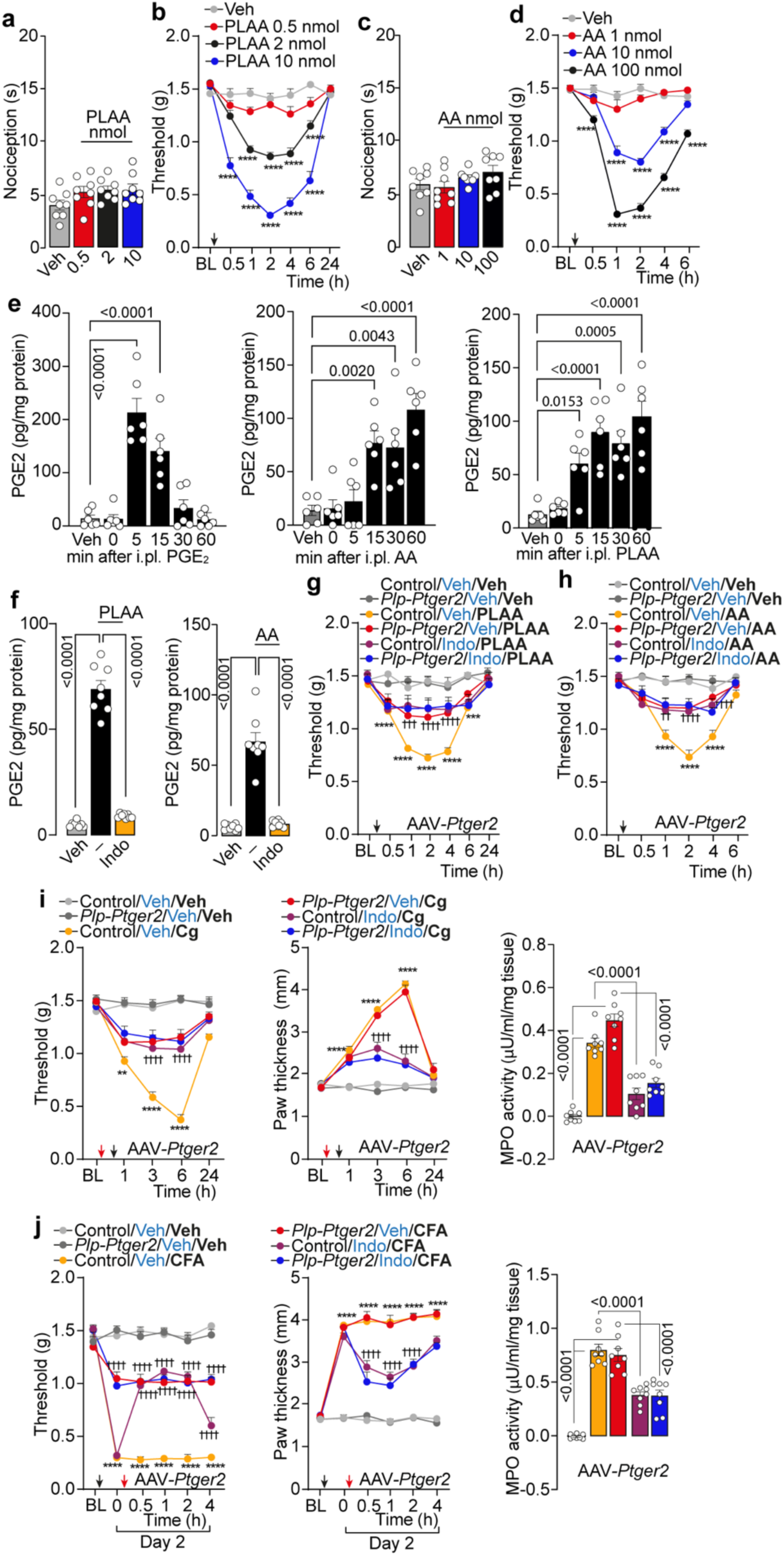
EP2 in Schwann cells mediates carrageenan and CFA hindpaw mechanical allodynia (allodynia) but not inflammation. **a, c**, Dose-dependent acute nociception and **b, d**, dose- and time-dependent allodynia after intraplantar (i.pl.) phospholipase A_2_ activating protein (PLAA), arachidonic acid (AA) or vehicle (Veh) in C57BL/6J mice (B6) (n =8 mice per group). **e,** Time dependent PGE_2_ assay in B6 paw tissue homogenates after i.pl. PGE_2_ (1.5 nmol), PLAA (2 nmol), AA (10 nmol) or Veh (n=6 mice per group) and **f,** pretreated with indomethacin (Indo, 280 nmol) (n=8 mice per group). **g, h,** Allodynia after i.pl. PLAA, AA or Veh in *P1p-Cre* or Control mice infected with AAV for selective silencing of EP2 (*P1p-Ptger2*) and pretreated with Indo or Veh (n =8 mice per group). **i, j**, Allodynia, paw thickness and MPO activity assay after i.pl. carrageenan (Cg, 300 μg), complete Freund’s adjuvant (CFA) or Veh in *Plp-Ptger2* or Control mice and pretreated with Indo or Veh (n =8 mice per group). Data are mean ± s.e.m. **a, c, e, f** and MPO activity in **i, j**, 1-way or **b, d, g, h,** allodynia and paw thickness in **i, j** 2-way ANOVA, Bonferroni correction. ***P<0.001, ****P<0.0001 vs. Veh or Control/Veh/Veh ^††^P<0.01, ^†††^P<0.001, ^††††^P<0.0001 vs. Control/Veh/PLAA, AA, Cg, CF A.

Next, we investigated two preclinical models of inflammatory pain, carrageenan and CFA, that evoke PG-dependent mechanical allodynia^30^. In control mice, i.pl. carrageenan or CFA increased PGE_2_ levels that were abated by indomethacin and did not produce detectable acute or delayed (assessed 3 h and 2 days after stimulus, respectively) non-evoked nociception (Supplementary Fig.7a,b). EP2 but not EP4 antagonism reduced the robust and sustained allodynia produced by carrageenan- or CFA similarly in male and female mice (Supplementary Fig.7c,d). An EP1 antagonist also failed to reduce mechanical allodynia by carrageenan and CFA (Supplementary Fig.7e). Allodynia evoked by both carrageenan and CFA was reduced in mice with EP2 silencing in SCs (*Plp-Ptger2* and *Plp-Ptger2-bis* mice) (Figure 2i,j and Supplementary Fig.7f), whereas no attenuation was detected in any other silenced mouse strains (Supplementary Fig.7g-m).

Importantly, allodynia was similarly inhibited in mice treated with indomethacin and in *Plp-Ptger2* mice, and indomethacin did not further increase the attenuation produced by SC EP2 silencing (Figure 2i,j). However, whereas indomethacin inhibited carrageenan- and CFA-induced increases in a series of inflammatory responses, including paw oedema and leucocyte myeloperoxidase (MPO), these responses were unchanged in *Plp-Ptger2* (Figure 2i,j) or in any other of the DRG neuron or SC silenced mouse strains (Supplementary Fig.7h-m). Pharmacological studies corroborated these findings as EP2 (PF), but not EP4 (BGC) antagonism, reduced carrageenan- or CFA-induced allodynia (Supplementary Fig.8a-d). Neither EP2 nor EP4 antagonism reduced carrageenan- and CFA evoked increase in additional inflammatory responses, including interleukin 1-β (IL-1β) and tumor necrosis factor-α (TNFα) (Supplementary Fig.8e,f). Carrageenan or CFA also increased a non-evoked pain-like response (grimace behavior) that was similarly attenuated by indomethacin, EP2 antagonism, and in EP2 *Plp-Ptger2* and *Plp-Ptger2-bis* mice, but not in *Plp-scr-shRNA* mice or EP4 antagonism (Supplementary Fig.8g). Intraplantar PGE_2_ (1.5 nmol/paw) failed to produce grimace behavior (Supplementary Fig.8h). These results in prototypical inflammatory pain models highlight the unexpected yet critical role of SC EP2 in PG-mediated non-evoked and evoked pain-like responses, which is totally independent from the PG-mediated inflammatory response.

### cAMP nanodomains in SCs signal pain

cAMP is a primary intracellular mediator of PG-induced inflammation and pain^31^. EP2 and EP4 couple to Gαs, which activates adenylyl cyclase (AC)^8^. cAMP and downstream cAMP-dependent PKA constitute a temporally and spatially organized nanodomain that controls different compartmentalized signaling pathways^32^. We recently reported that CGRP-stimulated periorbital mechanical allodynia is mediated by a sustained cAMP increase in SCs, which requires clathrin- and dynamin-dependent internalization and endosomal signaling of the CGRP-CLR/RAMP1 complex^15^. In contrast, we report here that clathrin and dynamin inhibitors (Pitstop2 and Dyngo-4a, respectively) failed to reduce PGE_2_, carrageenan- and CFA-evoked allodynia (Supplementary Fig.9). Thus, EP2 and cAMP signaling at endosomes of SCs is unlikely to mediate allodynia produced by inflammation.

To probe the contributions of compartmentalized cAMP signaling in SCs to EP2-dependent allodynia, we used blue light *Beggiatoa* photo-activable adenyl cyclase (bPAC)^33,34^, which enables optogenetic stimulation of focal cAMP production. Human SCs (hSCs) were co-transfected with bPAC and a genetically-encoded red fluorescent global cAMP sensor, cADDIs, to monitor cAMP production. Blue light (pulsed 1 s/5 s for 10 min) stimulated a forskolin-like increase in cADDIs fluorescence (ΔF/F0) peaking at 60 s and returning to baseline within 10 min (Figure 3a), as previously reported^35^. To study this SC signaling pathway *in vivo,* we used a *Plp^Cre^* driver that functions as a lineage tracer to express bPAC-T2A-mCherry in SCs. To induce the bPAC-selective expression in SCs, a Cre-dependent virus packaged with the AAV2/8 serotype was used for efficient infection of SCs. A loxP flanked bPAC-T2A-mCherry construct was used to exclusively express bPAC and mCherry in SCs (Figure 3b). Intraplantar virus injection in *Plp^Cre^*mice labelled cutaneous SCs (Figure 3c). Blue-light evoked cAMP-dependent allodynia in *Plp^Cre^* mice but not in control mice (Figure 3d). These data confirm the proalgesic role of cutaneous SCs^17^ which contribute to mechanical allodynia.

**Figure 3.**
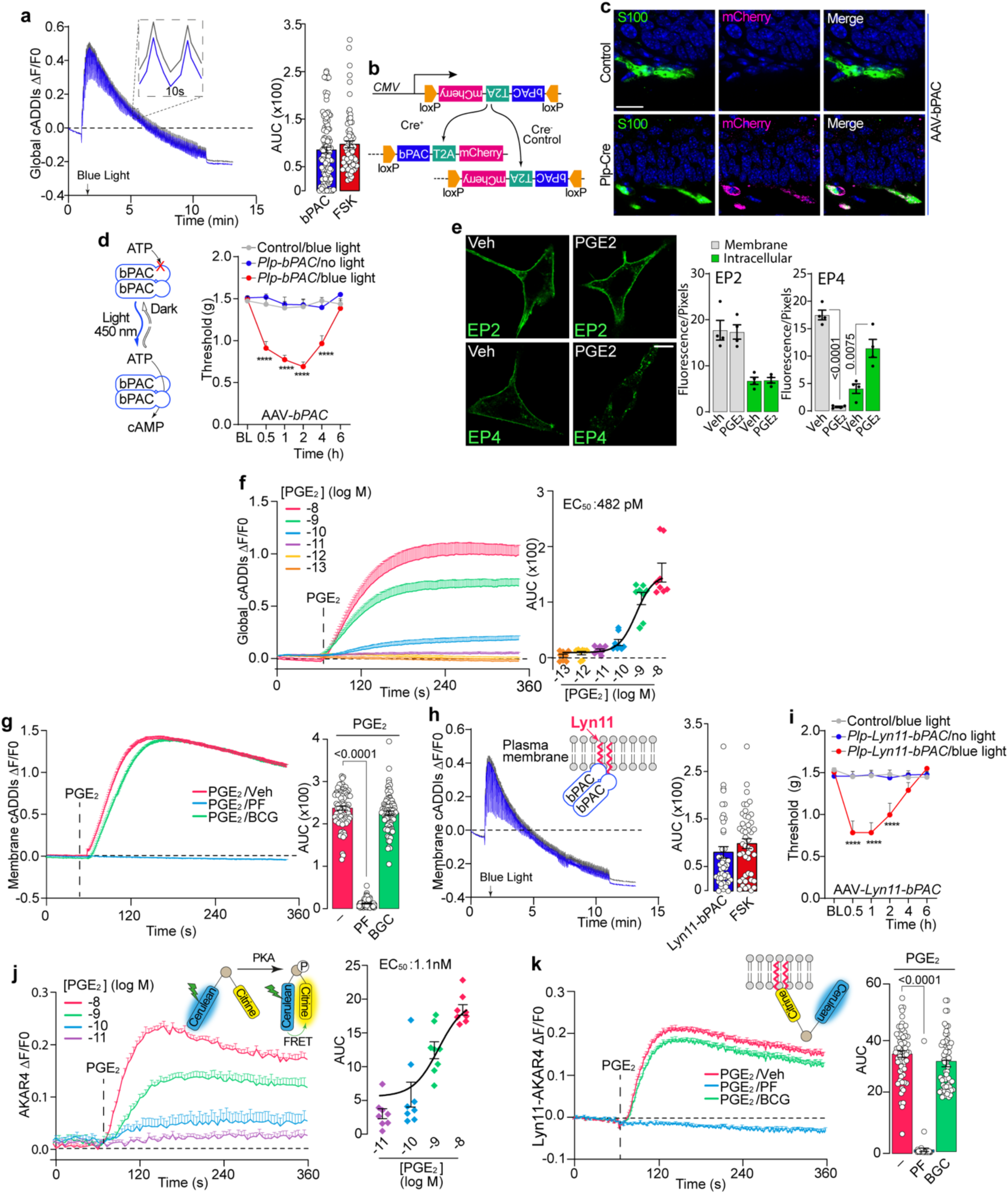
Schwann cells cAMP nanodomains promote hindpaw mechanical allodynia (allodynia) by PGE_2_. **a**, Global cAMP formation in human Schwann cells (hSCs), after blue light stimulation (450 nm, 10 min) (n=121 cells, n=3 independent experiments) or forskolin (FSK 100 μM) (n=78 cells, n=3 independent experiments). **b**, Schematic representation of *Cre* recombinase dependent expression of bPAC and mCherry separated by T2A self-cleaving peptide sequence. **c**, Representative images of mCherry expression in S100B+ cells in mouse paw tissue after intraplantar (i.pl.) infection with AAV for selective expression of bPAC in SCs (scale bar: 20 μm) (n=4 subjects). **d**, Illustration of bPAC activation after blue light stimulation and allodynia induced by blue light stimulation (pulsed 1 s/5 s for 10 min) in *Plp-Cre* or Control mice infected with AAV-*bPAC* (*Plp-bPAC*) (n=8 mice per group). **e**, Representative images and cumulative data of membrane and intracellular localization of EP2 and EP4 in hSCs cells, with or without pre-incubation with unlabeled PGE_2_ (10 µM, 30 min). Scale bar, 20 µm. (n=4 independent experiments). **f**, Concentration-dependent global cAMP formation in hSCs induced by PGE_2_ (n=8 replicates). **g**, Membrane confined cAMP formation induced by PGE_2_ (10 nM) in hSCs in the presence of PF-04418948 (PF,1 μM), BGC 20-1531 (BGC, 1 μM) or vehicle (Veh) (cells number: PGE_2_=80, PF=90, BGC=88, n=3 independent experiments). **h**, Illustration of membrane tagged Lyn11-bPAC membrane confined cAMP formation and cAMP formation in hSCs after blue light stimulation (n=48 cells, n=3 independent experiments) or FSK (100 μM) (n=46 cells, n=3 independent experiments). **i**, Allodynia induced by blue light stimulation in *Plp-Cre* or Control mice infected with AAV-*Lyn11-bPAC* (*Plp-Lyn11-bPAC*) (n =8 mice per group). **j**, Concentration-dependent PKA activation induced by PGE_2_ in hSCs (n=8 replicates). **k**, Membrane confined PKA activation in hSCs induced by PGE_2_ (10 nM) in the presence of PF (1μM), BGC (1 μM) or Veh (cells number: PF=64, BGC=63, PGE_2_= 64, n=3 independent experiments). Data are mean ± s.e.m. **a, e**, **h,** Student’s t test**, g, k** 1-way or **d, i** 2-way ANOVA, Bonferroni correction. AUC, area under curve. ****P<0.0001 vs. Control/blue light.

### SC plasma membrane-bound PKA/AKAP drives EP2 sustained pain signal

Observations in transfected HeLa cells show that PGE_2_ induces EP4 but not EP2 internalization^36^, suggesting that EP2-induced cAMP signaling in SCs is confined to the plasma membrane. We confirmed these findings in HEK 293T cells and hSCs expressing EP2 or EP4 C-terminally fused to Venus. PGE_2_ (10 µM) stimulated robust endocytosis of EP4 after 30 min, whereas EP2 remained at the plasma membrane (Figure 3e and Supplementary Fig.10a).

To identify the cAMP subcellular compartment implicated in allodynia, we studied total and nanodomain-specific cAMP signaling in SCs transfected with global cellular or plasma membrane-targeted cADDIs sensor. PGE_2_ concentration- and time-dependently increased global cADDIS fluorescence in hSCs that was attenuated by selective EP2 (PF) and EP4 (BGC) antagonism (Figure 3f and Supplementary Fig.10b, c). Increases in plasma membrane-associated cAMP evoked by PGE_2_ in hSCs, and in mouse SCs from sciatic nerve trunks or cutaneous nerve fibers, were attenuated by EP2 but not EP4 antagonism (Figure 3g and Supplementary Fig.10d,e). Peri-sciatic PGE_2_ failed to cause non-evoked nociception but elicited sustained allodynia measured in the hind paw, ipsilateral to the injection, which was attenuated by EP2 antagonism and in EP2 *Plp-Ptger2* and *Plp-Ptger2-bis* mice, but not by EP4 antagonism or in *Plp-scr-shRNA* mice (Supplementary Fig.10 f-k). Thus, *in vivo* experiments support the hypothesis that all SCs, both cutaneous and those wrapping more proximal sciatic nerve trunks, sustain allodynia.

The selective EP2 (butaprost) and EP4 (L-902) agonists increased global cAMP but only butaprost stimulated plasma membrane-associated cAMP (Supplementary Fig.10l-n). To determine the contribution of plasma membrane cAMP signaling to allodynia, we fused bPAC with the membrane targeting sequence, Lyn11 (Lyn11-bPAC). Plasma membrane localization of bPAC was confirmed in HEK293 cells (Supplementary Fig.10o). hSCs were cotransfected with Lyn11-bPAC and the plasma membrane-targeted cADDIS sensor (Figure 3h). Blue light increased plasma membrane cADDIs fluorescence similarly to forskolin, which was unaffected by EP2 or EP4 antagonism (Supplementary Fig.10p). Paw exposure to blue light in *Plp^Cre^* mice infected with i.pl. AAV-Lyn11-bPAC (*Plp-Lyn11-bPAC* mice) time-dependently enhanced allodynia (Figure 3i). Thus, a plasma membrane-associated cAMP nanodomain mediates allodynia.

cAMP and cAMP-dependent PKA constitute a spatiotemporally organized circuit that is strictly compartmentalized to form signaling nanodomains that control distinct biochemical and functional pathways^32^. The role of PKA in PGE_2_ and inflammatory mechanical hypersensitivity has been previously documented^31^. However, to the best of our knowledge, increases in membrane-associated cAMP and the ensuing intracellular pathway have not been previously associated with pain signaling. To determine if membrane-associated cAMP activates discrete compartmentalized complexes through spatially localized sequestered PKA, hSCs were transfected with global (AKAR4) or plasma membrane-targeted (Lyn-AKAR4) fluorescent resonance energy transfer (FRET)-based PKA activity biosensors. PGE_2_, butaprost (EP2 agonist) and L-902 (EP4 agonist) induced a concentration- and time-dependent increase in AKAR4 FRET signals (Figure 3k, Supplementary Fig.11a,b). Global PKA activation by PGE_2_ was similarly prevented by both EP2 (PF) and EP4 (BGC) antagonists (Supplementary Fig.11c). However, PF but not BGC inhibited PGE_2_-stimulated plasma membrane-associated PKA activity detected with the Lyn-AKAR4 biosensor (Figure 3l). Selective EP2 and EP4 activation confirmed these results; butaprost but not L-902 stimulated plasma membrane PKA activity detected with Lyn-AKAR4 (Supplementary Fig.11d).

To examine whether the SC plasma membrane-localized cAMP/PKA activity mediates PGE_2_-induced allodynia, we used a light-activated phosphodiesterase (LAPD)^37,38^ fused with the plasma membrane targeted sequence Lyn11 (Lyn11-LAPD). hSCs were co-transfected with Lyn11-LAPD and the membrane targeted cAMP FRET sensor Lyn11-R-FlincA. To confirm the plasma membrane localization of LAPD and R-FlincA HEK293 cells were used (Supplementary Fig.11e,f). PGE_2_ increased the FRET signal that was prevented by the red-light activation of the Lyn11-LAPD (Figure 4a). These findings were recapitulated *in vivo* as paw stimulation of *Plp^Cre^* mice, infected with i.pl. AAV(Lyn11-LAPD) (*Plp-Lyn11-LAPD* mice), with red-light prevented allodynia evoked by PGE_2_ (Supplementary Fig.11g). Exposure to red-light of *Plp-Lyn-LAPD* mice also reduced allodynia evoked by AA, PLAA, carrageenan and CFA (Figure 4b,c, Supplementary Fig.11h,i). These results confirm the essential role of plasma membrane-associated cAMP/PKA nanodomains in SCs in mediating mechanical allodynia induced by PGE_2_ and inflammation.

**Figure 4.**
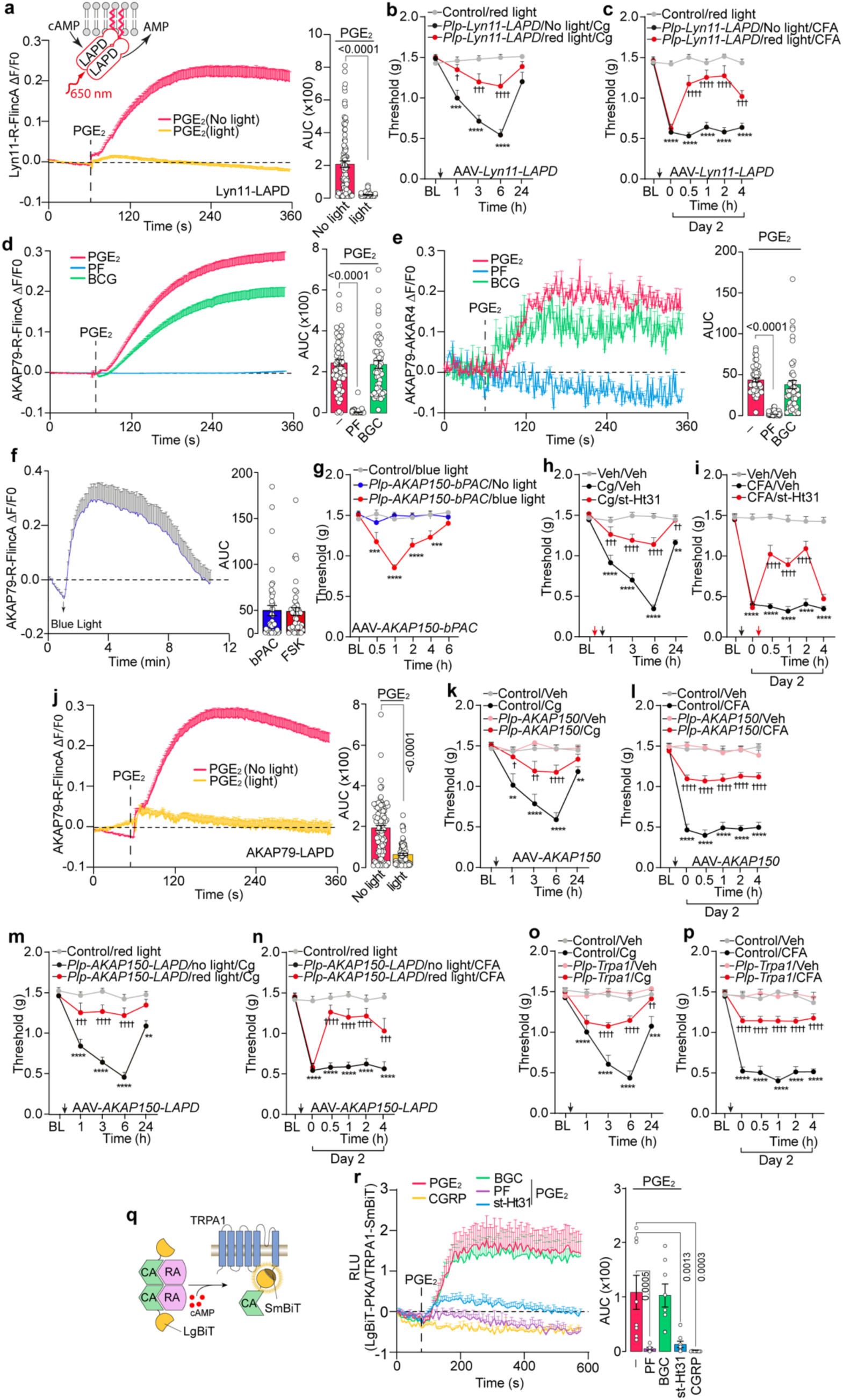
Schwann cells membrane-bound PKA/AKAP drives hindpaw mechanical allodynia (allodynia) by PGE_2_ and inflammation. **a**, Membrane cAMP formation in human Schwann cells (hSCs) after red light (650 nm, 10 min) activation of membrane-tagged light-activated phosphodiesterase (Lyn11-LAPD) and PGE_2_ (10 nM) (cells number: light=61, No light=97). **b, c,** allodynia induced by carrageenan (Cg, 300 μg) or complete Freund’s adjuvant (CFA) in *Plp-Cre* or Control mice after intraplantar (i.pl.) infection with AAV-*Lyn11-LAPD* (*Plp-Lyn11-LAPD*) and stimulated with red light (n=8 mice per group). **d,** AKAP79 associated cAMP formation and **e**, PKA activation induced by PGE_2_ (10 nM) in the presence of PF-04418948 (PF, 1 μM), BGC 20-1531 (BGC, 1μM) or vehicle (Veh) in hSCs (**d,** cells number: PGE_2_=62, PF=98, BGC=57, n=3 independent experiments; **e,** cells number: PF=95, BGC=47, PGE_2_=52, n=3 independent experiments). **f,** AKAP79 associated cAMP formation in hSCs after blue light stimulation (1s/5s) (n=43 cells, n=3 independent experiments) or forskolin (FSK, 100 μM) (n=46 cells, n=3 independent experiments). Allodynia induced by (**g)** blue light stimulation (1s/5s) in *Plp-Cre* or Control mice after i.pl. infection with AAV-*AKAP79-bPAC* (*Plp-AKAP79-bPAC*) (n=8 mice per group); **h**, **i**, Cg, CFA or Veh in C57BL/6J mice pretreated with st-Ht31 (17 nmol) or Veh (n=8 mice per group). **j**, AKAP79 associated cAMP formation in hSCs after red light activation of AKAP79-LAPD and PGE_2_ (10 nM) (cells number: light=102, No light=100, n=3 independent experiments). Allodynia induced by Cg, CFA or Veh in (**k**, **l**) *Plp-Cre* or Control mice infected with AAV for selective silencing of *AKAP150* (*Plp-AKAP150*) (n=8 mice per group); **m, n,** *Plp-Cre* or Control mice after i.pl. AAV-*AKAP79-LAPD* (*Plp- AKAP150-LAPD*) and stimulated with red light (n=8 mice per group); **o, p,** *Plp-Trpa1* or Control mice (n=8 mice per group). **q,** Illustration of BRET assay. **r**, Catalytic subunit of PKA (CA) and TRPA1 interaction after PGE_2_ (100 nM) in the presence of BGC (100 nM) PF(100 nM), st-Ht31 (10 μM) or after CGRP (10 μM) (replicates number: PGE_2_=5, BGC=5, PF=7, st-Ht31=7, CGRP=16). Data are mean ± s.e.m. **a, f, j** Student’s t test **d, e, r,** 1-way or **b, c, g, h, i, k-p,** 2-way ANOVA, Bonferroni correction. AUC, area under the curve. **P<0.01, ***P<0.001, ****P<0.0001 vs. Veh, Control/Veh, blue light, red light ^†^P<0.05, ^††^P<0.01, ^†††^P<0.001, ^††††^P<0.0001 vs. Cg, CFA, PGE_2_/Veh, Control/Cg, CFA, Plp-AKAP150-LAPD/no light/Cg, CFA.

In excitable cells, the plasma membrane-localized scaffold protein, AKAP79/150, tightly controls the phosphorylation of pain signaling proteins^7,39^. Due to its multivalent nature, we hypothesized that AKAP79/150 scaffolds cAMP/PKA at the plasma membrane of SCs and thereby spatiotemporally regulates PGE_2_-evoked allodynia. To determine whether AKAP79/150 recruits signaling effectors to distinct compartments of SCs that fine-tune PGE_2_ responses, we fused the cAMP FRET sensor R-FlincA or the PKA FRET sensor AKAR4 to the full-length AKAP79 scaffold (gene *AKAP5*). PGE_2_ induced a time-dependent increase in AKAP79-R-FlincA fluorescence in hSCs that was prevented by EP2 (PF) but not EP4 (BGC) antagonism (Figure 4d). Similarly, PGE_2_ stimulated AKAR4-AKAP79 fluorescence that was attenuated by EP2 but not EP4 antagonism (Figure 4e). The EP2 agonist, butaprost, but not the EP4 agonist, L-902, increased the AKAP79-R-FlincA fluorescence and AKAR4-AKAP79 FRET signals in hSCs, further supporting the critical contribution of the AKAP79 compartment in SC EP2 signaling (Supplementary Fig.12a,b).

To determine the contribution of the SC AKAP79/150-associated cAMP increase to allodynia, we fused bPAC to full-length AKAP79 (AKAP79-bPAC). In hSCs coexpressing AKAP79-bPAC and the AKAP79-R-FlincA FRET sensor, blue light stimulation increased FRET signal (Figure 4f). To translate these findings *in vivo*, blue light was applied to the paw of *Plp^Cre^* mice infected with i.pl. AAV AKAP150-bPAC (*Plp-AKAP150-bPAC* mice), leading to time-dependent allodynia (Figure 4g). To determine the contribution of AKAP79/150 scaffold to inflammatory pain *in vivo*, we used the peptide inhibitor of the AKAP/PKA interaction, st-Ht31^40^. st-Ht31 pretreatment inhibited carrageenan- or CFA-evoked allodynia in B6 mice (Figure 4h,i). In hSCs coexpressing the cAMP FRET sensor AKAP79-R-FlincA and AKAP79-LAPD, red-light activation of the AKAP79-LAPD prevented PGE_2_-stimulated cAMP production at the AKAP79 compartment (Figure 4j). Selective silencing of SC AKAP150 in *Plp^Cre+^* mice infected with intrasciatic AAV AKAP150 (*Plp-AKAP150* mice) reduced allodynia evoked by carrageenan and CFA (Figure 4k,l). Furthermore, paw stimulation of *Plp-AKAP150-LAPD* mice with red-light attenuated allodynia evoked by carrageenan or CFA (Figure 4m,n), underlining a mechanistic role of the SC PKA/AKAP interaction in inflammatory mechanical allodynia.

### EP2 associates with TRPA1 in SCs to signal pain

Transient receptor potential ankyrin 1 (TRPA1) is a key mediator of CGRP-evoked activation of SCs and mechanical allodynia. CGRP signals from endosomes of SCs to activate cAMP and PKA, resulting in endothelial nitric oxide synthase (eNOS) phosphorylation and nitric oxide (NO) release^15^. NO activates TRPA1, leading to formation of reactive oxygen species (ROS). ROS target TRPA1 on adjacent nociceptors to signal mechanical allodynia^15^. We investigated whether plasma membrane-associated cAMP/PKA nanodomains in SCs similarly promote mechanical hypersensitivity.

NOS inhibition attenuates PGE_2_-evoked mechanical hyperalgesia in rats^41^. We confirmed that the unselective NOS inhibitor, Nω-nitro-L-arginine methyl ester hydrochloride (L-NAME), partially reduced PGE_2_-induced allodynia in mice (Supplementary Fig.13a). The CGRP receptor antagonist, olcegepant, and the combination of L-NAME and olcegepant produced a similar inhibition of PGE_2_-induced allodynia (Supplementary Fig.13a), suggesting that a component of allodynia evoked by PGE_2_ is due to CGRP release from peptidergic nerve terminals and the ensuing NO release. In contrast to the data obtained with PGE_2_, L-NAME or olcegepant failed to reduce AA-, PLAA-, carrageenan- or CFA-evoked allodynia (Supplementary Fig.13b-i). Thus, endogenous PGs released during inflammation are unable to target peptidergic nerve terminals and release CGRP. However, selective TRPA1 silencing in SCs (*Plp*-*Trpa1* mice) markedly inhibited PGE_2_-evoked allodynia (Figure 4o,p and Supplementary Fig.13j). Despite the finding that endogenous PGs and CGRP activate signaling pathways that converge on TRPA1 as a common final pathway to signal allodynia, the following results indicate that the underlying mechanism for channel activation elicited by these mediators is different.

PKA anchoring to AKAP79/150 regulates hypersensitivity to inflammatory mediators, including sensitization of the TRP vanilloid 1 (TRPV1) channel by PGE_2_^7^. We hypothesized that a direct PKA/TRPA1 phosphoactivation in SCs elicits the L-NAME-independent allodynia in response to EP2 stimulation. A time-resolved NanoLuc complementation reporter approach was used to measure the dynamics of PKA/TRPA1 interaction in hSCs. Large (LgBiT) and small (SmBiT) subunits of the Nanoluc luciferase were fused to the C- and N-terminal extremities of TRPA1 and PKA catalytic subunit, respectively (SmBiT-TRPA1, PKA-LgBiT) (Figure 4q). PGE_2_ stimulated an increased luminescence in HEK293T cells coexpressing SmBiT-TRPA1 and PKA-LgBiT with EP2 but not with EP4 (Supplementary Fig.13k). In hSCs, PGE_2_, but not CGRP, increased luminescence. Antagonism of EP2 (PF) but not the EP4 (BGC) blocked this response (Figure 4r). Disruption of AKAP79/PKA interaction with the inhibitory peptide, st-Ht31, also attenuated PGE2-stimulated luminescence (Figure 4r). Thus, a first component of PGE_2_-induced allodynia results from a direct PKA/TRPA1 interaction while a second component depends on CGRP release and the ensuing NOS-induced NO release. In agreement with these data, allodynia by the EP2 agonist, butaprost, was completely attenuated by st-Ht31 and in *Plp*-*Trpa1* mice but was unchanged in mice treated with the CGRP receptor antagonist, olcegepant (Supplementary Fig.13l-n).

Several findings suggests that SC PKA/TRPA1 phosphoactivation by EP2/cAMP sustains calcium-dependent oxidative stress signal, which activates neuronal TRPA1 to evoke pain. PGE_2_ elicited a calcium-response and H_2_O_2_ generation; PF (EP2 antagonist) and A96 (TRPA1 antagonist) blocked these responses (Supplementary Fig.14a,b). Selective silencing of TRPA1 in DRG neurons (*Adv-Trpa1* mice) attenuated PGE_2_-evoked allodynia compared to control mice. However, the TRPA1-independent acute nociception to PGE_2_^42^ was similar in B6 and *Adv-Trpa1* mice (Supplementary Fig.14c,d). Allodynia produced by carrageenan or CFA was also attenuated in *Adv-Trpa1* (Supplementary Fig.14e,f) compared to control mice. The spin-trap agent, N-tert-butyl-alpha-phenylnitrone (PBN), which captures free-radicals, reduced allodynia to PGE_2_, carrageenan or CFA, but did not affect acute nociception to PGE_2_ (Supplementary Fig.14g-j). The observation that pretreatment but not post-treatment (30 min after stimulus) with the AC inhibitor, SQ22536, reduced PGE_2_-evoked allodynia (Supplementary Fig.14k,l) indicates that membrane-associated cAMP increase initiates but does not sustain mechanical hypersensitivity.

### SC EP2 blockade abates pain but does not delay pain resolution

Although NSAIDs transiently inhibit inflammatory pain, they markedly delay its resolution^43^. This delayed resolution of pain, reported in the CFA mouse model of inflammatory pain, and recapitulated in patients, is ascribed to disruption of the protective role of the PG-mediated inflammation that favors the recovery from the painful condition^43^. We observed that daily treatment (intraperitoneal, for 6 days after CFA) with the non-selective COX inhibitor, diclofenac, attenuated CFA-evoked allodynia as well as the increases in paw oedema and MPO levels (Figure 5a-c). After cessation of diclofenac treatment (at day 7 after CFA), allodynia persisted, and recovery was not attained within the following 25 days (Figure 5a). Although identical treatment with the EP2 antagonist, PF, also initially inhibited allodynia (like diclofenac), PF neither decreased inflammation (paw oedema, MPO levels) nor delayed allodynia recovery (Figure 5a-c). Furthermore, selective silencing of SC EP2 (*Plp-Ptger2*) in mice produced an early and sustained attenuation of CFA-evoked allodynia without reducing inflammation (Figure 5d-f). The observation that, like in *Plp-Ptger2* mice, in SC AKAP150 silenced (*Plp-AKAP150*) mice CFA-induced allodynia was constantly attenuated without any reduction in inflammation (Figure 5g-i) supports the unique role of the EP2-dependent intracellular pathway in SCs in the control of inflammatory allodynia.

**Figure 5.**
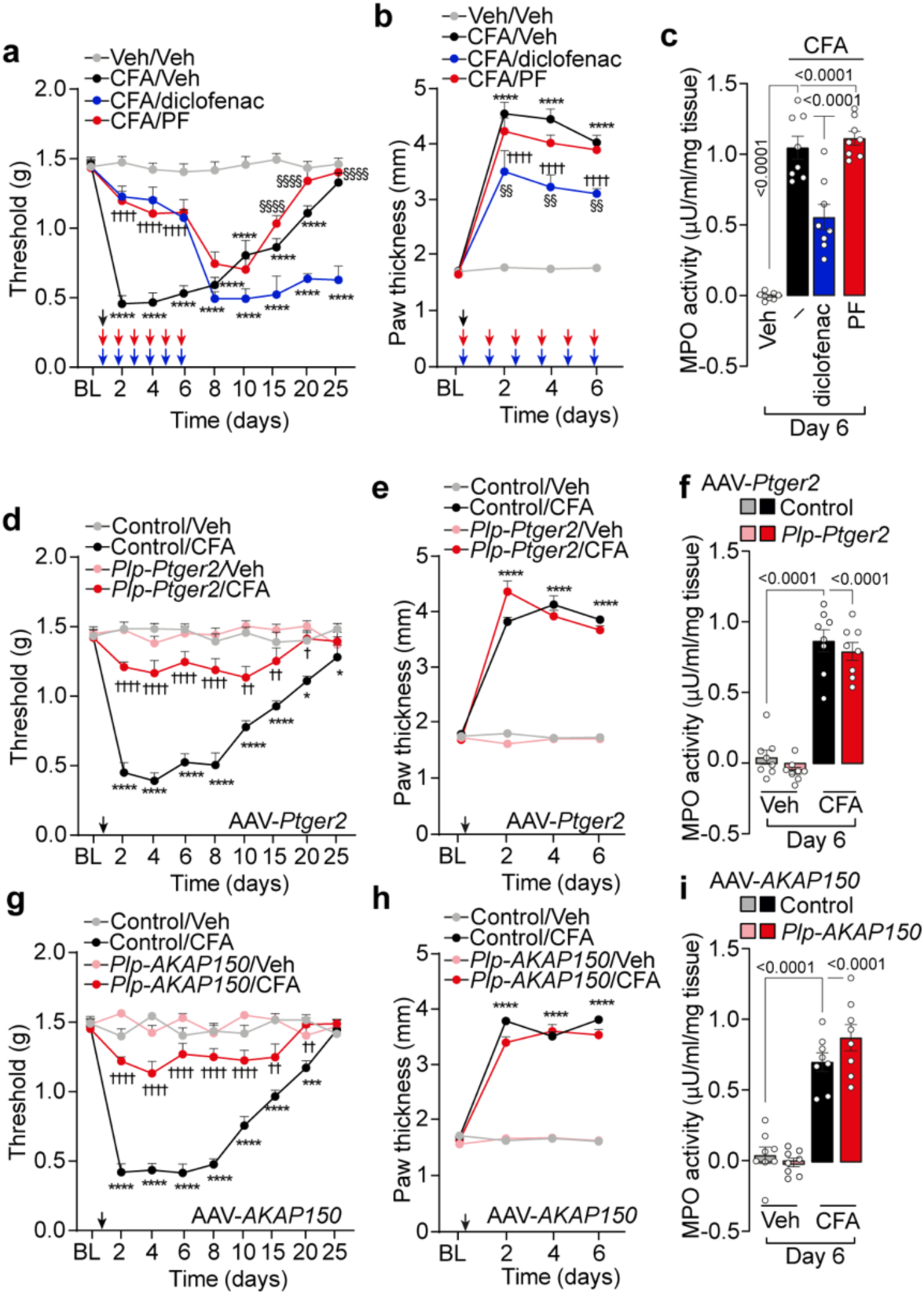
EP2 in Schwann cells blockade abates hindpaw mechanical allodynia (allodynia) but does not delay pain resolution. **a-c,** Allodynia, paw thickness and myeloperoxidase (MPO) activity assay after complete Freund’s adjuvant (CFA) or vehicle (Veh) in C57BL/6J mice treated for 6 consecutive days with diclofenac (25 mg/kg, i.p.), PF-04448948 (PF, 10 mg/kg, i.g.) or Veh (n=8 mice per group). **d-i** allodynia, paw thickness and MPO activity assay after CFA or Veh in *P1p-Cre* or Control mice infected with AAV for selective silencing of **d-f,** EP2 *(-Ptger2*) (*Plp-Ptger2*) or **g-i**, *AKAP150* (*Plp-Akap150*) (n=8 mice per group). Data are mean ± s.e.m. **a, b, d, e, g, h,** 2-way or **c,f, i,** 1-way ANOVA, Bonferroni correction. *P<0.05, ***P<0.001, ****P<0.0001 vs. Veh, Control/Veh ^†^P<0.05, ^††^P<0.01, ^††††^P<0.0001 vs. CFA/Veh, Control/CFA, ^§§^P<0.01, ^§§§§^P<0.0001 vs. CFA/diclofenac.

## Discussion

Since discovery of the mechanism of action of aspirin^44^, the analgesic effect of NSAIDs has been intimately associated with their prominent anti-inflammatory activity. Herein, we challenge the dogma that for relief of PG-mediated inflammatory pain it is necessary to block PG-induced inflammation. In fact, we provide pharmacological and genetic evidence that PGs released during inflammation act locally to stimulate two mechanistically independent pathways, one of which mediated by SC EP2 produces pain-like behaviors without any implication in the inflammatory response. The observations that SCs taken from cutaneous nerve fibers or sciatic nerve trunks responded similarly to EP2 agonism, antagonism, and silencing, and that hind paw allodynia produced by either intraplantar or peri-sciatic PGE_2_ injection was mechanistically similar, suggest that the EP2 proalgesic role is not limited to SCs localized to cutaneous terminal nerve fibers. The SC proalgesic pathway initiated by EP2 results in TRPA1-dependent ROS release that targets neuronal TRPA1, leading to sustained hypersensitivity to mechanical stimuli (Figure 6).

**Figure 6.**
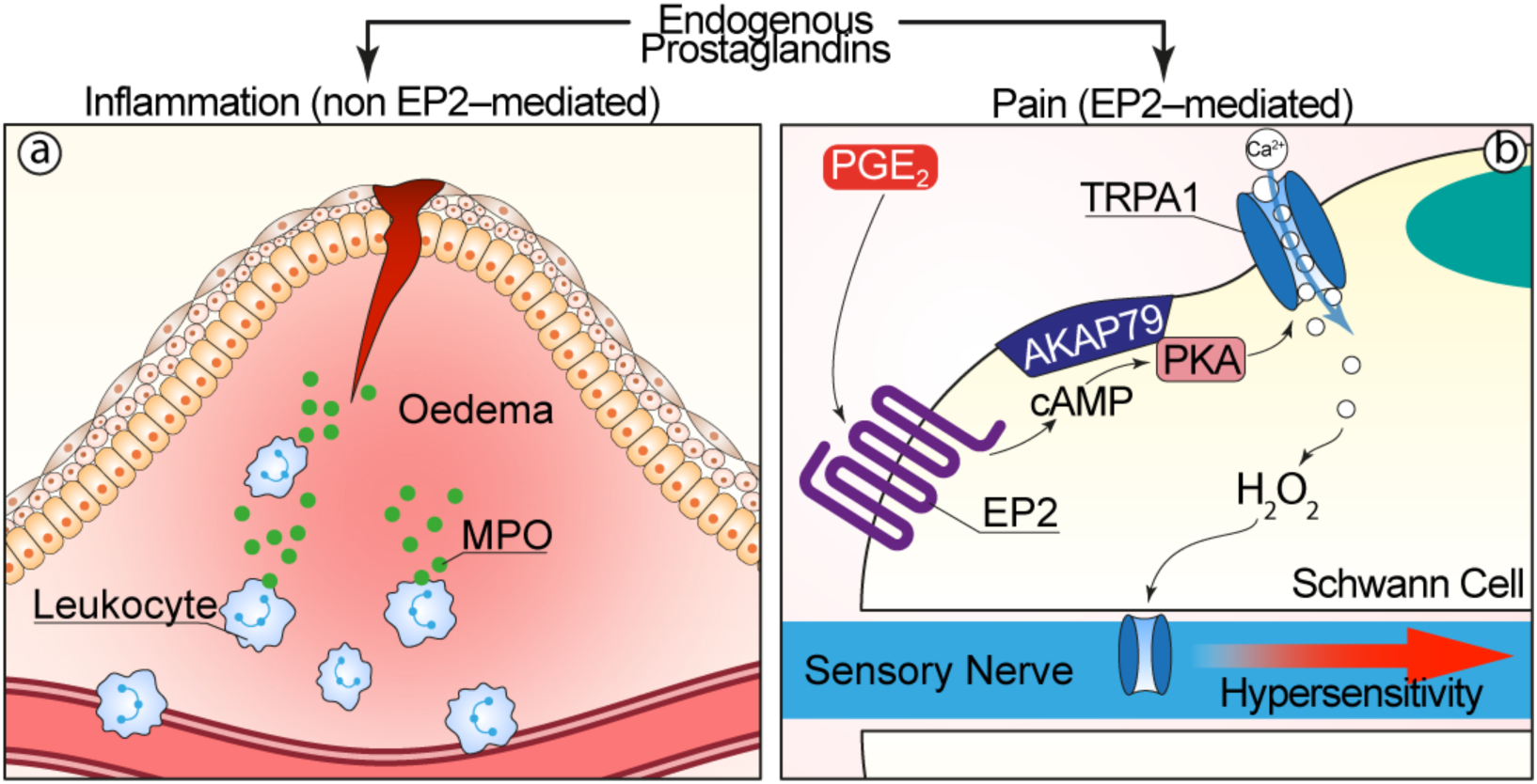
Hypothesized mechanism of action of PGs. PGs released during inflammation promote inflammation and pain by distinct mechanisms. **a,** PGs cause inflammation by an EP2-independent inflammatory process, which is characterized by oedema, leukocyte infiltration and release of myeloperoxidase (MPO). **b,** PGs cause pain by activating EP2 on SCs. EP2 generates cAMP, leading to AKAP79-associated activation of PKA in plasma membrane-delimited nanodomains. PKA phosphorylates and activates TRPA1 in SCs, which elicits a calcium-dependent release of reactive oxygen species (H_2_O_2_). H_2_O_2_ targets TRPA1 on adjacent nociceptors, which results in sustained hypersensitivity to mechanical stimuli.

Our results exclude the role of neuronal EP4 in mediating inflammatory pain hypersensitivity^10^ and that the proalgesic action of EP2 is confined to the central nervous system^45^. Some previous studies implicated EP2 in pain signaling in the periphery, including reports that EP2 deletion attenuates mechanical allodynia evoked by intradermal PGE_2_ or zymosan^45^, and that the selective EP2 antagonist, PF, attenuates primary peripheral hyperalgesia in an endometriosis mouse model^46^. Scant attention has been paid, however, to the existence of PGE_2_ receptors in peripheral glial cells and to their contribution to pain. EP1 and EP4 expression was reported in myelinated SCs^27^, and EP2 was documented in SC dental pulp^47^ and oligodendrocytes^28^.

Our findings do not support the currently accepted view that the PGE_2_ injection in peripheral tissues recapitulates the painful response elicited by endogenous PGs, since we found that the actions of exogenous and endogenous PGs differ significantly. On one hand, we confirm that PGE_2_ injection elicits both a robust, acute and transient non-evoked nociception and a moderate delayed allodynia by directly targeting EP4 in DRG nociceptive nerve terminals. On the other hand, we reveal that SC EP2 mediates both evoked (mechanical allodynia) and non-evoked (grimace behavior) pain-like responses elicited by endogenous PGs, released under inflammatory circumstances. It is possible that the higher PGE_2_ tissue level achieved locally after the injection of the exogenous compound may more likely target the dense network of sensory nerve terminals in the mouse hind paw. The observation^48^ that subcutaneous injection of a small dose of PGE_2_ in human skin elicits long-lasting hyperalgesia but not acute spontaneous pain is consistent with the present findings and may be due to the lower density of cutaneous nerve fibers in human as compared to mouse skin^49^ and the increased probability for injected PGE_2_ to target sensory nerve terminals in mice *vs.* men.

The unique role of SC EP2 to signal pain-like responses is highlighted by the receptor-mediated engagement of membrane cAMP nanodomains and membrane-associated AKAP/PKA to elicit pain-like responses *in vivo*. Our results expand the repertoire of SC intracellular mediators implicated in pain signaling, adding to endosomal cAMP increased by CGRP^15^, the plasma membrane cAMP compartment increased by PGs as essential pain signals. A limitation of the present study is that currently available tools do not allow identification of the SC subtype, myelinated or Remak, mechanistically implicated in inflammatory mechanical allodynia and grimace behavior, an issue that should be investigated by future research. Other limitations are the biases intimately associated with translation from *in vitro* to *in vivo* data and between species from mice to humans.

Recent evidence has highlighted the dual and opposing roles of NSAIDs. Although NSAIDs provide immediate relief from inflammatory pain, they substantially delay its resolution^43^. We reproduced the delayed and insufficient allodynia recovery provoked by diclofenac in the CFA model of inflammatory pain. However, the EP2 (PF) antagonist, while attenuating allodynia, did not prevent its resolution. Notably, selective silencing of EP2 and one of its downstream intracellular signaling mechanisms (AKAP150) in SCs attenuated both the initial and the sustained phase of inflammatory allodynia, providing pain relief superior to all the other interventions. Together, our results highlight the unique role of SC EP2 in inflammatory mechanical hypersensitivity and the dissociation of this proalgesic pathway from the concomitant PG-mediated inflammatory response that appears necessary for a timely and efficient pain resolution. Thus, novel therapeutic strategies that selectively block SC EP2 would provide better and safer treatments for inflammatory pain in patients and particularly in the elderly.

## AUTHOR CONTRIBUTIONS

R.N., F.D.L., P.G., N.W.B conceived and conceptualized the study; L.L., G.D.S, E.B. performed in vivo experiments, M.M. R.T., I.S., M.T., C.J.P., performed in vitro imaging experiments; M.C., M.Mo., A.R, P.P., G.B. performed AAV design and generation; DSMdA, A.M., V.A., V.D.G., D.D.J. performed immunofluorescence, RNA Scope and invitro assays; R.N., F.D.L. P.G., N.W.B. wrote the original draft of the paper; R.N., F.D.L., P.G., N.W.B., B.L.S., R.T., J.Z., reviewed and edited the paper with input from all authors.

## ACKNOWLEDGMENTS

Supported by grants from: European Research Council (ERC) under the European Union’s Horizon 2020 research and innovation programme (grant agreement No. 835286) (P.G.), National Institutes of Health (NS102722, DE026806, DK118971, DE029951, N.W.B., B.L.S.; R01 DK073368, R35 CA197622, J.Z.), Department of Defense (W81XWH1810431, W81XWH2210239, N.W.B., B.L.S.), European Union - Next Generation EU, National Recovery and Resilience Plan, Mission 4 Component 2 - Investment 1.4 - National Center for Gene Therapy and Drugs based on RNA Technology - CUP B13C22001010001 (R.N.) and Mission 4 Component 2 - Investment 1.3 - Mnesys A multiscale integrated approach to the study of the nervous system in health and disease – CUP B83C22004910002 (P.G., F.D.L.). Views and opinions expressed are however those of the author(s) only and do not necessarily reflect those of the European Union or the European Commission. Neither the European Union nor the European Commission can be held responsible for them.

## Methods

### Animals

Male and female mice C57BL/6 J (Charles River, RRID: IMSR_JAX:000664) were used throughout (25–30 g, 6–8 weeks old). To generate mice in which the *Trpa1* gene was conditionally silenced in Schwann cells, homozygous 129S-Trpa1^tm2Kykw/J^ (floxed Trpa1, *Trpa1^fl/fl^*, RRID:IMSR_JAX: 008649 Jackson Laboratory) were crossed with hemizygous B6.Cg-Tg(Plp1-CreERT)3Pop/J mice (*Plp-Cre^ERT^*, RRID: IMSR_JAX:005975 Jackson Laboratory) expressing a tamoxifen-inducible Cre in Schwann cells (Plp1, proteolipid protein myelin 1)^12^. The progeny (*Plp-Cre^ERT^;Trpa1^fl/fl^*) was genotyped using PCR for *Trpa1* and *Plp-Cre^ERT^.* Mice that were negative for *Plp-Cre^ERT^* (*Plp-Cre^ERT–^;Trpa1^fl/fl^*) were used as control. Both positive and negative mice for *Cre^ERT^* and homozygous floxed Trpa1 (*Plp-Trpa1* and control respectively) were treated with intraperitoneal (i.p.) 4-hydroxytamoxifen (4-OHT, 1 mg/100 μL in corn oil once a day consecutively for 3 days). To selectively delete *Trpa1* in primary sensory neurons, Trpa1^fl/fl^ mice were crossed with hemizygous Advillin-Cre mice (Adv-Cre)^12,50^. Mice positive or negative for Cre and homozygous for floxed Trpa1 (*Adv-Trpa1* and control respectively) were used. The successful Cre-driven deletion of TRPA1 mRNA was confirmed using reverse transcription quantitative real-time PCR (RT-qPCR). Some *Plp-Cre^ERT^*^+^ or *Plp-Cre^ERT^*^-^ were treated with 4-OHT (i.p., 1 mg/100 μL in corn oil, once a day consecutively for 3 days) before the infection with AAV for selective silencing of the different genes in Schwann cells. Hemizygous Advillin-Cre mice (*Adv-Cre^+^)* or their control (*Adv-Cre^-^*)^12^ were also used for the infection with AAV for selective silencing of the different genes in primary sensory neurons. The group size of n=8 mice for behavioral experiments was determined by sample size estimation using G Power [v3.1^51^] to detect the size effect in a *post-hoc* test with type 1 and 2 error rates of 5% and 20%, respectively. Allocation concealment of mice into the vehicle(s) or treatment groups was performed using a randomization procedure (http://www.randomizer.org/). The assessors were blinded to the identity of the animals (genetic background) or allocation to treatment groups. None of the animals were excluded from the study. Mice were housed in a temperature- and humidity-controlled *vivarium* (12 h dark/light cycle, free access to food and water, 5 animals per cage). At least 1 h before behavioral experiments, mice were acclimatized to the experimental room and behavior was evaluated between 9:00 am and 5:00 pm. Behavioral studies followed Animal Research: Reporting of *In Vivo* Experiments (ARRIVE) guidelines^52^. Animal experiments and sample collections were carried out according to the European Union (EU) guidelines for animal care procedures and Italian legislation (DLgs 26/2014) application of the EU Directive 2010/63/EU. All animal studies were approved by the Animal Ethics Committee of the University of Florence and the Italian Ministry of Health (permits no. 765/2019-PR, 288/2021-PR). Animals were anesthetized with a mixture of ketamine and xylazine (90 mg/kg and 3 mg/kg, respectively, i.p.) and euthanized with inhaled CO_2_ plus 10-50% O_2_.

## Behavioral experiments

### Treatment protocols

Butaprost and L-902.688 were purchased from MedChemExpress LLC. If not otherwise indicated, the reagents were obtained from Merck Life Science SRL or Tocris Bioscience. Mice received intraplantar (i.pl., 10 μl/site) injection of PGE_2_ (0.1, 0.5, 1.5, 5 nmol) or its vehicle (1% ethanol in 0.9% NaCl); (R)-Butaprost (0.1, 0.5, 1.5, 5 nmol), L-902,688 (0.5, 1.5 and 5 nmol), arachidonic acid (1, 10, and 100 nmol) or their vehicle (4% dimethyl sulfoxide, DMSO, 4% Tween 80 in 0.9% NaCl); phospholipase A_2_ activating protein (0.5, 2, and 10 nmol), λ-carrageenan (300 μg), complete Freund’s adjuvant (CFA, 1 mg/ml) or their vehicle (0.9% NaCl). PF-04448948 (5 nmol), BGC 20-1531 (1, 5, 10 nmol), indomethacin (280 nmol), SC-51322 (5 nmol), Ro-1138452 (5 nmol), L-798,106 (5 nmol), L-NAME (1 μmole), ER-819762 (1, 5, 10 nmol), Dyngo-4a (500 pmol), PitStop2 (500 pmol) and SQ25536 (25 nmol) or their vehicle (4% DMSO, 4% Tween80 in 0.9% NaCl), olgecepant (1 nmol) and st-Ht31 (17 nmol) or their vehicle (1% DMSO in 0.9% NaCl), PBN (1 μmol) or its vehicle (0.9% NaCl) were administered (10 μL, i.pl.) 30 min before the algogenic stimuli (i.pl.). In the CFA model, drugs were administered at day 2 after CFA injection. In another set of experiment, diclofenac (25 mg/kg, i.p.) or PF-04448948 (10 mg/kg, intragastric, i.g.) or their vehicle (4% DMSO, 4% Tween80 in 0.9% NaCl and 0.5% carboxymethyl cellulose, respectively) were administered once a day from day 0 to day 6 after CFA injection. Perisciatic injection was made as previously reported^53^. Briefly, PGE_2_ (1.5 nmol) or veh were injected in the region surrounding the sciatic nerve at high thigh level of the right hind limb (from ∼ 10 to ∼16 mm from the paw surface) without skin incision in a volume of 6 μl using a microsyringe with a 30-gauge needle. The position of sciatic nerve at high thigh level was chosen by using the femoral head as a landmark. *Plp-Cre^+^* and control or *Adv-Cre^+^* and control mice were infected with an intrasciatic (i.sc., 5 µL, 1×10^12^ v/g) or intrathecal (i.th., 5 µL,1×10^12^ v/g) injection of different AAVs. For intrasciatic injection mice were anesthetized and sciatic nerve was exposed. A volume of 3 µl of viral vectors was directly injected into the sciatic nerve through a 33-gauge needle and a Hamilton syringe connected to a manual micropump (World Precision Instruments). The needle remained in place at the injection site for 1 additional min, before it was slowly removed. Animals were used 3 weeks after AAVs infection. Sciatic nerve and DRGs were harvested for evaluating AAVs infection. In optogenetics experiments *Plp-Cre*^+^ and control mice infected with an intraplantar (i.pl., 10 µL, 1×10^12^ v/g) injection of different AAVs were used. After 3 weeks, the hind paw of infected mice was stimulated with a blue light (pulsed, 1 s/4 s, 450-nm light, 30 LED intensity) or red light (not pulsed, 650-nm light, 48 LED intensity) for 10 min before i.pl. CFA, Cg, or vehicle. The hind paw was harvested for evaluating AAV infection.

## Behavioral assays

### Acute nociception

Immediately after i.pl. injection, mice were placed inside a plexiglass chamber, and acute nociception response was assessed for 15 min by measuring the time (sec) that the animal spent in lifting, biting, licking, shaking the injected paw. The acute nociceptive response was also assessed in mice 3 h and 2 days after i.pl. Cg, CFA, respectively or vehicle.

### Mechanical allodynia

The mechanical paw-withdrawal threshold was measured using von Frey filaments of increasing stiffness (0.02–2 g) applied to the plantar surface of the mouse hind paw, according to the up-and-down paradigm^54^. The 50% mechanical paw-withdrawal threshold (g) response was then calculated from the resulting scores. Mechanical paw-withdrawal threshold was measured at baseline and at different time following treatments.

### Paw thickness

Paw thickness (in millimeters) was measured using 0-12 mm stainless digital thickness gauge (Mitutoyo Corporation). Paw thickness was measured at baseline, and at different time following i.pl. Cg, CFA or vehicle and indomethacin or vehicle.

### Grimace score

Spontaneous pain was tested using the Mouse Grimace Scale^55^. Briefly, each mouse was placed in an individual chamber (9×5×5 cm) having transparent Plexiglas walls to allow experimental observation. Mice were habituated for 30 min before behavioral testing. Cameras directed at the front of the cubicle recorded 30 min of facial expressions. One clear facial image was taken for every 3 min interval, scrambled and scored blindly for facial grimacing^55^. The scorers assessed facial expressions such as orbital tightening (closing or narrowing of the eyelid and orbital area), ear position (outward or backwards rotation of the ears), scoring either a 0 (not present), 1 (moderately visible) or 2 (severe) depending on the magnitude of the expression. Nose and cheek bulging, and whisker change were not score because they were difficult to distinguish against the dark fur of the mice. The test was performed in mice 3 h and 2 days after i.pl. Cg, CFA, respectively or vehicle.

#### Primary human Schwann cells

Primary human Schwann cells (hSCs) (#P10351, Innoprot or #10HU-188, iXCells Biotechnologies) were cultured and maintained in Schwann cell medium (#P60123, Innoprot or #MD0055, iXCells Biotechnologies) at 37 °C in 5% CO_2_ and 95% O_2_ as previously reported^13^. After 12 passages, cells were discarded and replaced.

#### HEK293T cell line

Human kidney epithelial (HEK293T) cells (#CRL-3216™, American Type Culture Collection) were cultured in DMEM supplemented with FBS (10%), L-glutamine (2 mM), penicillin (100 U/ml) and streptomycin (100 mg/ml) at 37 °C in 5% CO_2_ and 95% O_2_.

#### AAVpro293T cell line

AAVpro 293T cells (#632273, Takara), were maintained in DMEM high glucose supplemented with 10% heat inactivated FBS, 4 mM L-glutamine,1 mM penicillin/streptomycin and 1 mM sodium pyruvate at 37 °C in 5% CO_2_ and 95% O_2_. The day before transfection, cells were plated in DMEM supplemented with 2% FBS.

#### Sciatic nerve primary mouse Schwann cells

Primary culture of mouse Schwann cells (mSCs) was obtained from sciatic nerves of C57BL/6J mice^15^. Briefly, the *epineurium* was removed, and nerve explants were cut (1 mm segments) and dissociated in Hank’s Balanced Salt Solution (HBSS, 2 h, 37 °C) using hyaluronidase (0.1%) and collagenase (0.05%). After centrifugation (150xg, 10 min, RT), the pellet was resuspended and cultured in DMEM containing streptomycin (100 mg/ml), fetal calf serum (10%), forskolin (2 μM), L-glutamine (2 mM), penicillin (100 U/ml) and neuregulin (10 nM). Cytosine arabinoside (Ara-C, 10 mM) was added three days later, to remove fibroblasts. Cells were cultured at 37 °C in 5% CO_2_ and 95% O_2_ for 15 d before experiments by replacing the culture medium every 3 days.

#### Cutaneous primary mouse Schwann cells

Mouse cutaneous Schwann cells were isolate as reported ^56^ ^57^ (with some modifications. Briefly, mouse hindpaws were collected on ice cold HBSS, washed in HBSS and fat and blood vessels were gently removed from skin under a light microscope. This skin was cut into small cubes and digested in dispase (5 mg/ml) for 2 h at 37°C in agitation (200 rpm). Epidermis was discarded and the dermis was minced and digested in collagenase Type IV (2 mg/ml) for 2 h at 37°C in agitation (200 rpm). The tissues were centrifuged and washed twice in HBSS to remove debris and then dissociated by mechanical pipetting in 5 ml of HBSS containing DNase enzyme (400 µg/ml), and further incubated for 1 h at 37°C in agitation (200 rpm). Samples were then centrifuged (300 xg) for 5 min and the pellet resuspended and cultured in DMEM supplemented with FBS (10 %), forskolin (2 μM), L-glutamine (2 mM), penicillin (100 U/ml), N2 supplement (1%). Cytosine arabinoside (Ara-C, 10 mM) was added three days later, to remove fibroblasts. Cells were cultured at 37 °C in 5% CO_2_ and 95% O_2_ for 6 days.

#### Constructs

Using NanoBiT® MCS Starter System (#N2014, Promega), N- and C-terminal LgBiT and SmBiT fusions proteins were generated. Eight different constructs amplifying PKA and hTRPA1 synthetized by Vector Builder: pBit2.1-C PKA-SmBit (P1/P2 and NheI-HF/SacI); pBit2.1-N-SmBIT-PKA (P3/P4 with SacI/XbaI); pBit1.1-C-hTRPA1-LgBIT (P5/P6 with AsiSI/XhoI); pBit1.1-N-LgBIT- hTRPA1 (P7/P8 with XhoI/XbaI) were generated according to manufacturer’s instructions. Using a Gibson assembly strategy, pBit1.1-C-PKA-LgBit (P9/P10 for PKA fragment and P11/P12 for acceptor vector) and pBit2.1-N-SmBit-hTRPA1 (P13/P14 for hTRPA1 fragment and P15/P16 for acceptor vector) were cloned. pBit1.1-N-LgBit-PKA was generated by Gibson assembly strategy with P17/P18 for PKA fragment and P19/P20 for acceptor vector. pBit2.1-C-hTRPA1-SmBit was generated by amplifying hTRPA1 and cloned in pBit2.1 acceptor vector, P21/P22 and P23/P24 primers were used. Plasmids were cloned using Gibson Assembly Cloning Kit (#E5510S, New England Biolabs). pBit1.1-N-LgBit-PKA and pBit2.1-C-hTRPA1-SmBit was the only combination with a BRET signal after PGE_2_ stimulation.

### In vitro studies

pCDNA3-AKAR4, pCDNA3-AKAP79-AKAR4, pCDNA3-Lyn11-AKAR4 and pCDNA3-AKAP79-(Ci-Ce)-Epac2 were used^32^. A blue light photo-activable adenyl cyclase from the *Soil Bacterium Beggiatoa* (bPAC) was amplified from pAAV-CMV-bPAC-T2A-mCherry (Vector Builder) and cloned into pCDNA3. Primers P25/P26 and NheI-HF/EcoRI-HF were used. pCDNA3-Lyn11-Lyn11-bPAC was generated by cloning the Lyn11-Lyn11 sequence into the pCDNA3-bPAC plasmid, P27/P28/P29/P30 oligos and NheI-HF/BamHI were used. pCDNA3-AKAP79-bPAC was generated by amplifying AKAP79 from pCDNA3-AKAP79-AKAR4^32^ and cloned into pCDNA3-bPAC, P31/P32 primers and NheI-HF/BamHI were used. pCDNA4-AKAP79-R-FlincA was generated by replacing AKAP79 from pCDNA3-AKAP79-(Ci-Ce)-Epac2 in pCDNA4-R-FlincA (<u>R</u>ed <u>Fl</u>uorescent <u>in</u>dicator for <u>cA</u>MP), kindly donated by Dr. Horikawa K^58^), BamHI-HF/XhoI were used. pCDNA4-Lyn11-Lyn11-R-FlincA was generated by replacing AKAP79 in the pCDNA4-AKAP79-R-FlincA with Lyn11-Lyn11 expressing adaptors P33/P34/P35/P36; sticky ends from HindIII-HF/BamHI were used. pCDNA3-LAPD was generated by replacing bPAC in the pCDNA3-bPAC with LAPD from pCG084 plasmid, gently donated by Dr. Möglich A^37^, P37/P38 primers and NheI-HF/XbaI were used. pCDNA-AKAP79-LAPD was generated by cloning AKAP79 from pCDNA3-AKAP79-bPAC into pCDNA3-LAPD, using NheI sticky ends. P39/P40 primers were used. pCDNA3- Lyn11-Lyn11-LAPD was generated by annealed Lyn11-Lyn11 expressing adaptors (P41/P42/P43/P44) and cloned with NheI-HF.

### In vivo studies

pAAV EF1-alpha-MCS was generated by replacing the mCherry in the pAAV-EF1-alpha rev(mCherry) with a multicloning site (MCS), P45/P46 primers were used. EF1-alpha-promoter was replaced with a short-CMV to obtain pAAV-CMV-MCS that was used as acceptor plasmid for all the constructs. pAAV-CMV-bPAC-T2A-mCherry was generated by cloning loxP-bPAC-T2A-mCherry-loxP from pAAV-CAG-bPAC-T2A-mCherry (Vector Builder) into pAAV-CMV-MCS, P49/P50 primers and SpeI-HF/HpaI were used. pAAV-CMV-LAPD-FLAG and pAAV-CMV-FLAG-LAPD were generated by PCR amplification from pCG084 of LAPD-FLAG and FLAG-LAPD and cloning into pAAV-CMV-MCS, P51/P52 or P53/P54 primers and SpeI-HF/HpaI and NheI-HF/SnaBI were used respectively. pAAV-CMV-Lyn11-Lyn11-bPAC-T2A-mCherry and pAAV-CMV-Lyn11-Lyn11-LAPD-FLAG were generated by cloning a Lyn11Lyn11 expressing adaptors (P33/P34/P35/P36) into pAAV-CMV-bPAC-T2A-mCherry and pAAV CMV-LAPD-FLAG, AvrII/HpaI were used. pAAV-CMV-AKAP79-bPAC-T2A-mCherry and pAAV-CMV-AKAP79-LAPD-FLAG were generated by cloning AKAP79 from pCDNA3-AKAP79-AKAR4, into pAAV-CMV-bPAC-T2A-mCherry and pAAV-CMV-LAPD-FLAG, P31/P55 primers and NheI-HF/AvrII were used. To not exceed the cargo limit of AAVs, WPRE and BGH poly(A) were removed from pAAV-CMV-AKAP79-LAPD-FLAG (EcoRV/PmlI) and WPRE sequence was removed from pAAV-CMV-AKAP79-bPAC-T2A-mCherry (ClaI). All constructs were confirmed by Sanger sequencing. All primers (P) sequences are available in the Supplementary Table 1. Constructs pAAV[FlexOn]-CMV-EGFP-[shRNA]-mCherry, expressing mPtger2, mPtger4 and mAkap5 shRNAs, were obtained using the following: mPTGER2 shRNA; 5′-ACCTTGGGTCTTTGCCATACTTTAGTGAAGCCACAGATGTAAAGTATGGCAAAGACCCAAGGG-3′; mPTGER4 shRNA, 5′-ACCAGTGAAACTCTGAAATTATTAGTGAAGCCACAGATGTAATAATTTCAGAGTTTCACTGGG-3′; mAKAP5 shRNA, 5′- CCAGTATGAAACACTCTTAATATAGTGAAGCCACAGATGTATATTAAGAGTGTTTCATACTGT -3′ (VectorBuilder). To obtain plasmids coding for a second short hairpin RNA targeting mPtger2 or mPtger4 or scrambled shRNA as negative control, the plasmid pAAV[FLEXon]-CMV LL-rev(EGFP-5’ miR-30E-BfuAI ORF_44bp BfuAI-3’ miR-30E)-rev(LL)-WPRE was designed and ordered (Vectorbuilder). shRNAs targeting the gene of interest (P56/P57, mPtger2; P58/P59, mPtger4; Vectorbuilder) or scrambled shRNA (P60/P61 scrambled mPtger2; P62/P63, Invivogen) were cloned by replacing the 44bp ORF with preannealed and phosphorylated oligonucleotides by using BfuAI compatible bases.

#### AAV generation

Recombinant AAV particles (rAAVs) were produced by using triple transfection strategy as described previously^13^. In brief, AAVpro 293T cells (#632273, Takara), were transfected with polyethylenimine (#23966, PEI, Polyscience) with a DNA:PEI ratio of 1:3 ^59^. To obtain rAAVs, AAVpro 293T cells were transiently transfected with 2.5 mg total DNA (plasmid expressing genes of interest, pAdDeltaF6; #11287, Addgene and Rep/Cap, 1:1:1 molar ratio). To infect with high efficiency Schwann cells and primary sensor neurons, Rep/Cap 2/8 or 2/9n were used, respectively (pAAV2/8 #112864; pAAV 2/9n #112865, Addgene). rAAVs were extracted and isolated 72 h post-transfection, purified by iodixanol gradient ultracentrifugation, concentrated, and titrated using a RT-qPCR assay (#6233 AAVpro Titration Kit, Takara) according to the manufacturer instructions.

#### Live cell imaging

HSCs were plated in 96-well poly-L-lysine-coated (8.3 μM) black clear bottom plates (5 × 10^5^ cells/well; PerkinElmer) and transfected with cDNA (130-300 ng) using jetOPTIMUS® DNA transfection reagent (#55-250, Polyplus) for 16-24 h at 37 °C in 5% CO_2_ and 95% O_2_. The day of experiments, hSCs were washed and added with HBSS at pH 7.4 at 37 °C. All the experiments were performed using a fluorescent microscope for recording (Axio Observer 7; with a fast filter wheel and Digi-4 lens to record excitations and Ultra-fast Sutter Lambda DG4 Xenon excitation source (range 300-700 nM) (Zeiss) with 20x or 40x objectives or using FlexStation3 Multi-Mode Microplate Reader (Molecular Devices) with SoftMax® Pro7 software (Molecular Devices).

#### cAMP *in vitro* imaging

To measure cAMP signaling in real time, cells were infected with a cADDIs BacMam virus (1.09 × 10^9^ v/g/mL) encoding the green upward cAMP sensor (#U0200G, Montana Molecular). The fluorescent signal was recorded by microscopy ex/em 506/517 nm (filter set: FT 495, ex BP 470/40, em BP 525/50, interval 1 s) or using FlexStation3 (ex/em 485/515 nm, interval 1.5 s), a Red Membrane-Targeted fS15 cADDIs membrane sensor (#U0241R, Montana Molecular), ex/em 558/603 nm (filter set: FT 570, ex BP 550/25, em BP 605/70, interval 1 s) or a R-FlincA ex/em 568/592 nm (filter set: FT 495, ex BP 470/40, em BP 525/50, interval 1 s). HSCs transfected with pCDNA4-R-FlincA were cultured at 32 °C for 24 h before imaging. HSCs expressing the green upward cAMP sensor were stimulated with PGE_2_ (100 fM-10 nM). The response to PGE_2_ (100 pM) was evaluated in the presence of PF-04418948 (100 pM-1 µM) or BGC20-1531 (100 pM-1 µM). In another set of experiments, cells were stimulated with L-902,688 (1 nM-10 µM) or (R)-Butaprost (10 nM-100 µM). HSCs, mSCs from sciatic nerves and cutaneous derived SCs expressing Red Membrane-Targeted fS15 cADDIs membrane sensor, were stimulated with PGE_2_ (10 nM) in the presence of PF-04418948 (1 µM) or BGC20-1531 (1 µM). In hSCs, single concentration of L-902,688 (100 µM) or (R)-Butaprost (100 µM) were tested. With R-FlincA sensor, hSCs were stimulated with PGE_2_ (10 nM) in the presence of PF-04418948 (1 µM) or BGC20-1531 (1 µM). Single concentration of L-902,688 (100 µM) or (R)-Butaprost (100 µM) were tested. Signals were recorded for 360 s, the ΔF/F0 ratio was calculated for each experiment and the results were expressed as the area under the curve (AUC).

bPAC was used to selectively induce an optogenetic cAMP increase in different subcellular compartments. HSCs were co-transfected with pCDNA-bPAC or pCDNA- Lyn11-Lyn11-bPAC and the red fluorescent cAMP sensor cADDIs (#U0200R for total cAMP, or #U0241R for Lyn11-Lyn11, Montana Molecular), or with pCDNA-AKAP79-bPAC and the pCDNA-AKAP79-R-FlincA sensor. HSCs were stimulated with a 450 nm light (pulsed, 1s/5s, 30% led intensity) for 10 min (led, #UHP-T-450-EP, controller #UHPTLCC-02-USB, Pulser USB for TTL pulse train generator, Prizmatik) and recorded with the fluorescent microscope (ex/em 558/603 nm; filter set: FT 570, ex BP 550/25, em BP 605/70, acquisition every 1 s for total and Lyn11-Lyn11-bPAC and 5 s for AKAP79-bPAC). Forskolin (100 μM) was used as positive control for cAMP increase. In some experiments the membrane cAMP formation after activation of Lyn11-bPAC was evaluated in the presence of PF-04418948 (1 µM) or BGC 20-1531 (1 µM).The ΔF/F0 ratio was calculated for each experiment and the results were expressed as AUC define.

To further evaluate the role of cAMP, hSCs were co-transfected with pCDNA-Lyn11-Lyn11-LAPD and pCDNA-Lyn11-Lyn11-R-FlincA or, with pCDNA-AKAP79-LAPD (AKAP79/150-LAPD) and AKAP79/150-R-FlincA. HSCs were exposed to a 650 nm light continuously (48% light intensity) for 10 min (#UHP-T-650-EP, Prizmatikl) and then stimulated with PGE_2_ (10 nM).

#### PKA activity *in vitro* imaging

To evaluate PKA activity, the FRET-based PKA activity sensor AKAR4 (pCDNA3-AKAR4) was used (filter set: channel1 FT 455, ex BP 436/25, em BP 535/30, interval 1 s, channel2 BS 420, ex BP 436/25, em BP 480/40, interval 1 s). HSCs were stimulated with PGE_2_ (100 pM-10 nM). The response to PGE_2_ (10 nM) was evaluated in the presence of PF-04418948 (1 µM) or BGC 20-1531 (1 µM). HSCs were exposed to different concentrations of (R)-Butaprost (10 µM - 500 µM) or L-902,688 (10 µM-500 µM). HSCs transfected with plasma membrane associated AKAR4 (pCDNA3-Lyn-AKAR4) or plasma membrane-localized scaffold protein AKAP79 (pCDNA3-AKAP79- AKAR4) were stimulated with PGE_2_ (10 nM) in the presence of PF-04418948 (1 µM) or BGC 20-1531 (1 µM). HSCs were stimulated with single concentrations of (R)-Butaprost (100 µM) or L-902,688 (100 µM). FRET changes were measured as a ratio of the acceptor fluorophore emission (545 nm) to donor emission (480 nm). Signals were recorded for 360 s. The ΔF/F0 ratio was calculated for each experiment and the results were expressed as the AUC.

#### Calcium imaging

HSCs were plated on poly-L-lysine-coated (8.3 μM) 35 mm glass coverslips and maintained at 37 °C in 5% CO_2_ and 95% O_2_ for 24 h. Cells were loaded (40 min) with Fura-2 AM-ester (5 μM) added to the buffer solution (37 °C) containing (in mM) 2 CaCl_2_; 5.4 KCl; 0.4 MgSO_4_; 135 NaCl; 10 D-glucose; 10 HEPES and bovine serum albumin (BSA, 0.1%) at pH 7.4. Cells were washed and transferred to a chamber on the stage of a fluorescent microscope for recording (Olympus IX 81) and exposed to PGE_2_ (10 nM) or vehicle (0.9% NaCl) in the presence of PF (1 μM) or A96 (50 μM) and the calcium response was monitored. Results were expressed as percent increase in ratio340/380 over baseline normalized to the maximum effect induced by ionomycin (5 μM) added at the end of each experiment.

#### H_2_O_2_ *in vitro* imaging

A genetically encoded probe for H_2_O_2_-HyPer [HyPer7.2^60^, kindly donated by Dr. Emrah Eroglu, Harvard Medical School, Boston, US] was used on live cells. HSCs were plated on poly-L-lysine-coated (8.3 μM) 35-mm glass coverslips and transfected with DNA (2 μg) of HyPer7.2 using jetOPTIMUS^®^ DNA transfection reagent (#55-250; Polyplus, Lexington, MA, USA). After 24–48 h, the hSCs were washed and transferred to a chamber on the stage of a fluorescent microscope for recording (Axio Observer 7; with a fast filterwheel and Digi-4 lens to record excitations; ZEISS, Stuttgart, Germany). Cells were exposed to PGE_2_ (10 nM), or its vehicle (0.9% NaCl), in the presence of PF (1 μM) or A96 (50 μM) and H_2_O_2_ variations were monitored for approximately 15 min. Results were expressed as the percentage increase in the ratio_408/455_ over the baseline, the ΔF/F0 ratio was calculated for each experiment and the results were expressed as the AUC.

#### Nanobit complementation assay

To evaluate the dynamics of PKA-TRPA1 proximity in hSCs, a time-resolve luciferase re-complementation assay was used. HSCs were plated in 96-well poly-L-lysine-coated (8.3 μM) white clear bottom plates (5 ×10^5^ cells/well; #6005181, PerkinElmer) the day before transfection. pBit2.1-C-hTRPA1-SmBit was co-transfected with pBit1.1-N-LgBit-PKA (130 ng DNA) with jetOPTIMUS® DNA transfection reagent (#55-250, Polyplus, Lexington). Forty-eight h after the transfection cells were treated for 10 min with Nano-Glo® Live Cell Reagent, according to manufacture instructions (#N2011, Nano-Glo^®^ Live Cell Assay System Promega). Then, hSCs were stimulated with PGE_2_ (100 nM), CGRP (10 µM) or vehicle. The response to PGE_2_ (100 nM) was evaluated in the presence of PF-04418948 (100 nM) or BGC 20-1531 (100 nM) or st-Ht31 (10 μM) or their vehicle (0.001% DMSO). To evaluate the dynamic of PKA-TRPA1 proximity in HEK293T cells, the same approach of hSCs was used. SmBit-TRPA1 was co-transfected with PKA-LgBit (130 ng DNA) with Ptger2 or Ptger4 expressing plasmids (6.5bng of DNA) using PEI (#23966, Polyscience). Forty-eight h after the transfection HEK293T cells were stimulated with PGE_2_ (100 nM). Luminescence of both experiments was measured for 6 min using a luminescence plate reader (FlexStation 3 Multi-Mode Microplate Reader; Molecular Devices) with SoftMax^®^ Pro7 software (Molecular Devices). The results are expressed as arbitrary units (AU) The ΔF/F0 ratio was calculated for each experiment and the results were expressed as the AUC.

#### EP2-Venus and EP4-Venus trafficking

cDNA encoding human EP2 and EP4 were cloned with a C-terminal monomeric (m) Venus tag on the C-terminus with a flexible linker (LRPLGSSGGGGGGSG). HSCs transfected with EP2-Venus or EP4-Venus (3 µg/10 cm dish), were plated in poly-D-lysine-coated 35 mm glass-bottomed dishes. After 48 h, cells were washed in HBSS. PGE_2_ (10 µM) or Veh was added, and imaging obtained after 30 minutes incubation. Images were obtained using a Leica SPi8 confocal microscope (63X objective) and processed in ImageJ.

#### Immunofluorescence

Anesthetized mice were transcardially perfused with PBS and 4% paraformaldehyde. Lumbar DRGs and sciatic nerve tissues were collected, postfixed for 24 h, and paraffin-embedded or cryoprotected in 30% sucrose. Human DRG (#0062-HP-240, Gentaur), mouse DRG, human sciatic nerve (#0062-HP-261, Gentaur) and mouse sciatic nerve formalin-fixed paraffin-embedded (FFPE) sections (5 µm) were incubated with the following primary antibodies EP2 (#ab167171, rabbit monoclonal, Abcam, 1:250), EP4 (#BS-8538R, rabbit polyclonal, Bioss, 1:200), S100B (#MA1-26621, mouse monoclonal, Invitrogen, 1:50), NeuN (#MAB377, mouse monoclonal, Merck, 1:250), diluted in fresh blocking solution (PBS, pH 7.4, 5% normal goat serum (NGS)), after antibody retrieval preformed with tri-sodium citrate buffer (pH 6) for EP2 and tris-EDTA buffer (pH 9) for EP4. Sections were then incubated with the fluorescent polyclonal secondary antibodies Alexa Fluor® 488 (#A32731, goat polyclonal anti-rabbit, Invitrogen, 1:600) and Alexa Fluor® 594 (#A11005, goat polyclonal anti-mouse, Invitrogen, 1:600), and coverslipped using the mounting medium with DAPI (#ab104139, Abcam). Cryosections (10 µm) of mouse sciatic nerve tissues were incubated with the following primary antibodies: EP4 (#sc- 55596, mouse monoclonal, Santa Cruz, 1:50), Alexa Fluor® 488 Anti-S100B antibody (#ab196442, rabbit monoclonal, Abcam, 1:100), diluted in blocking solution 5% NGS. Sections were then incubated with the fluorescent polyclonal secondary antibodies Alexa Fluor® 594 (#A21207, donkey polyclonal anti rabbit, Invitrogen, 1:600) and Alexa Fluor® 594 (#A11005, goat polyclonal anti mouse, Invitrogen, 1:600), and coverslipped using the mounting medium with DAPI. All slides were visualized and analyzed using a Zeiss Axio Imager 2 microscope with Z-stacks in the Apotome mode (Zeiss).

#### Dual RNAScope Fluorescent in-situ hybridization and immunohistochemistry

Fluorescent *in situ* hybridization was performed using RNAscope™ Multiplex Fluorescent V2 Assay (ACDbio), according to manufacturer’s protocol. MSCs and hSCs were plated on fibronectin (#21370, SERVA) coated slides and cultured until confluence. Cells were fixed with paraformaldehyde 4% in PBS for 30 min at RT and dehydrated in 50%, 70% and 100% ethanol, for 1 min. Cells were rehydrated by subsequent steps in 70%, 50% ethanol and PBS followed by a permeabilization with PBS + 0.1% Tween 20 for 10 min at RT. Pretreatment was completed by adding 2-3 drops of H_2_O_2_, to inactivate endogenous peroxidases, followed by 2-3 drops of Protease III (ACDbio), diluted 1:15 in PBS for 10 min at RT. Slides were incubated at 40 °C in HybEZ II Oven (ACDbio). Cells were hybridized for 2 h with probes, to detect Ptger2 and Ptger4 in human (#406791, #406771, Biotechne) and mouse (#456481, #441461, Biotechne) cells. Negative control probe (#310043, Biotechne) was used as negative control. Signal amplification was obtained by consecutive incubations with AMP1, AMP2 and AMP3 reagents and HRP enzyme (ACDbio) for 15 min, followed by TSA Vivid fluorophore (#323271, ACDbio), diluted 1:1500 in TSA Buffer (#322809, ACDbio). Any further HRP activity was stopped by incubation with HRP blocker for 15 min. For immunofluorescence, hSCs cells were permeabilized in PBS + 0.1% Triton 100 for 15 min at RT. Cells were then blocked in in fresh blocking solution (PBS, pH 7.4, 5% normal donkey serum (NDS) and incubated O/N at 4 °C with primary anti-SOX10 antibody (#AF2864, goat polyclonal, R&D Systems, 1:200). Cells were than incubated with secondary antibodies Alexa Fluor® 594 (#A11058, donkey polyclonal anti goat, Invitrogen, 1:600) for 2 h at RT and coverslipped using the mounting medium with DAPI. All slides were visualized and analyzed using a Zeiss Axio Imager 2 microscope with Z-stacks in the Apotome mode (Zeiss). HEK293T cells were transfected with pCDNA Lyn11-Lyn11-LAPD (2 µg) and CAAX-rGFP (2µg) or Rab5a-LAPD (2µg) and Rab5a-Venus (2µg) using polyethylenimine (#23966, PEI, Polyscience) with a DNA:PEI ratio of 1:3, incubated overnight and plated on 12 mm glass coverslips. Cells were incubated overnight and fixed (4% paraformaldehyde in PBS) on ice for 20 min. Cells were washed 3 x in PBS and blocked in PBS + 0.3% saponin + 3% NHS for 1 h at RT. Cells were incubated with mouse monoclonal anti-6xHis antibody (1:1000, Takara) in PBS + 0.3% saponin + 1% NHS overnight at 4° C. Cells were washed and incubated with donkey anti-mouseAlexa568 (1:1000, Invitrogen) for 1 h at RT, counterstained with DAPI (1µg/ml), washed 3x in PBS and mounted with ProlongGlass hard set mounting medium (Thermofisher). Cells were imaged on a Leica SP8 confocal, and images were processed with ImageJ and Adobe Illustrator. HEK293T cells were transfected with pCDNA Lyn11-Lyn11-R-FlincA (0.2 µg) on 12 mm glass coverslips with polyethylenimine (#23966, PEI, Polyscience). After 24 h, transfected HEK293T cells were imaged on a Leica confocal microscopy (Stellaris 5), and images were processed with Adobe Illustrator.

#### Reverse transcription quantitative real-time PCR (RT-qPCR)

The total RNA from hSCs and mSCs or DRGs and sciatic nerve tissue of C57BL/6J, Plp-Cre and Adv-Cre mice was extracted by standard TRIzol method together with RNeasy Mini Kit (#74106, QIAGEN) according to manufacturer’s protocol, while total RNA of human DRGs was commercially purchased (#636150, Clontech). RNA concentration and purity were assessed spectrophotometrically by measuring the absorbance at 260/280 nm. The purified RNAs were retro-transcribed using SuperScript IV One-Step RT-PCR System (#12595025, Thermo Fisher) according to manufacturer’s protocol. For relative quantification of mRNA compared to housekeeping gene, real-time PCR was carried out using Rotor Gene Q (Qiagen Spa). The chosen reference gene was GAPDH. The KAPA SYBR FAST Universal Kit (#KK4602) was used for amplification, and the cycling conditions were as follows: samples were heated to 95 °C for 10 min followed by 40 cycles of 95 °C for 10 sec, and 65 °C 95 °C for 20 sec. PCR reaction was performed in triplicate for each sample. Relative expression of mRNA was calculated using the 2^−Δ(ΔCT)^ comparative method, with each gene normalized against the GAPDH gene for the same sample. The sets of primers are reported in Supplementary Table 2.

#### PGE_2_ IL-1β and TNF-α assay

PGE_2_ IL-1β and TNF-α content was measured in the hind paw homogenates using a single-analyte enzyme-linked immunosorbent assay kit (#ab287802, #ab197742, #ab208348Abcam) according to the manufacturer’s protocol. Data are expressed as pg/mg of protein.

#### Myeloperoxidase (MPO) activity assay

MPO activity was measured in the hind paw homogenates using a fluorometric assay kit (#ab111749, Abcam) according to the manufacturer’s protocol. Data are expressed as μU/ml/mg of tissue.

#### Statistical analysis

The results are expressed as the mean ± SEM. For multiple comparisons, a one-way ANOVA followed by a post-hoc Bonferroni’s test was used. The two groups were compared using Student’s *t*-test. For behavioral experiments with repeated measures, a two-way mixed-model ANOVA followed by a post-hoc Bonferroni’s test was used. Statistical analyses were performed on raw data using GraphPad Prism 8 (GraphPad Software Inc.). P*-*values less than 0.05 (*P* < 0.05) were considered significant. EC_50_ values were determined from non-linear regression models using Graph Pad Prism 8. The statistical tests used and sample size for each analysis are shown in the Fig. legends.

## Data availability

All materials generated in this study are available upon reasonable request from Francesco De Logu and Romina Nassini.

