## supplementary figures and tables for "Targeting the Schwann Cell EP2/cAMP Nanodomain to Block Pain but not Inflammation"

**Supplementary Figure Legends**

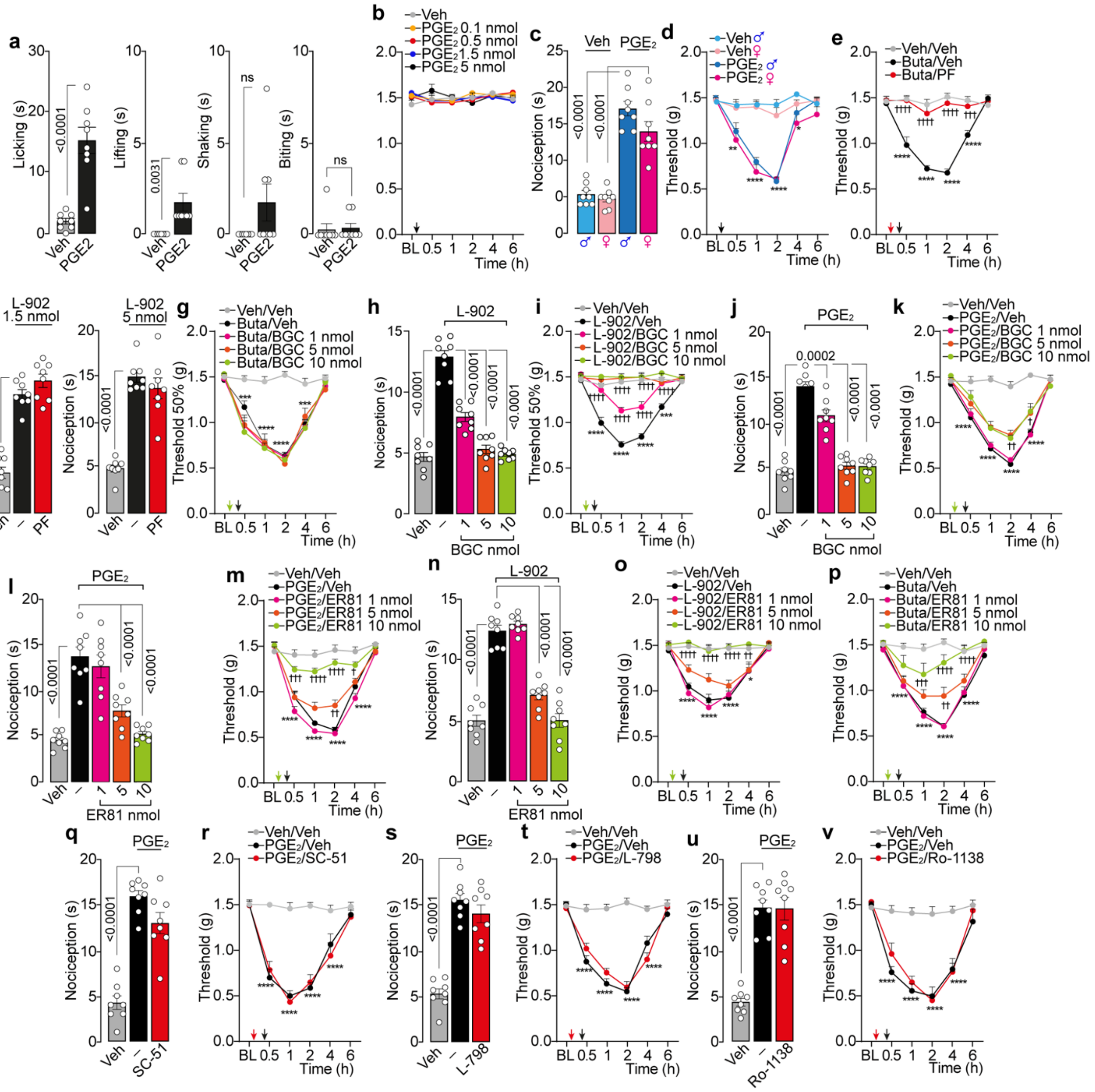

**Supplementary Fig. 1. EP2 Schwann cells mediates hind paw mechanical allodynia (allodynia) by PGE₂. a,** Licking, lifting, shaking, and biting behaviour after intraplantar (i.pl.) injection of PGE₂ (1.5 nmol) or vehicle (Veh) in C57BL/6J mice (B6). **b,** Allodynia in the contralateral paw after i.pl. PGE₂ (1.5 nmol) or Veh in B6 mice. **c, d,** Acute nociception, and allodynia in male and female B6 after i.pl. PGE₂ (1.5 nmol) or Veh. **e**, Allodynia after i.pl. (R)-Butaprost (Buta, 1.5 nmol) or Veh in B6 pretreated with PF-04448948 (PF, 5 nmol) or Veh. **f**, Acute nociception after i.pl. L-902,688 (L-902, 1.5 and 5 nmol) or Veh in B6 pretreated with PF (5 nmol) or Veh. **g**, Allodynia after i.pl. Buta (1.5 nmol) or Veh in B6 pretreated with BGC-20-1531 (BGC). Acute nociception and allodynia after i.pl. **h, i,** L-902 (5 nmol) or **j, k,** PGE₂ (1.5 nmol) or Veh in B6 pretreated with BGC or

Veh. Acute nociception and allodynia after i.pl. **l, m**, $PGE_2$ (1.5 nmol) or **n, o,** L-902 (5 nmol) or **p,** Buta (1.5 nmol) or Veh in B6 mice pretreated with ER-819762 (ER81) or Veh. **q, s, u,** Acute nociception and **r, t, v,** allodynia after i.pl. $PGE_2$ (1.5 nmol) or Veh in B6 mice pretreated with SC-51322 (SC-51, 5 nmol), L-798,106 (L-798, 5 nmol), Ro-1138452 (Ro-1138, 5 nmol) or Veh (all data, n=8 mice per group). Data are mean ± s.e.m. **a**, Student's t test, **c, f, h, j, l, n, q, s, u** 1-way or **b, d, e, g, I, k, m, o, p, r, t, v** 2-way ANOVA, Bonferroni correction. *P<0.05, **P<0.01, ***P<0.001, ****P<0.0001 vs. Veh $^{†}$P<0.05, $^{††}$P<0.01, $^{†††}$P<0.001, $^{††††}$P<0.0001 vs. Buta, L-902, $PGE_2$/Veh.

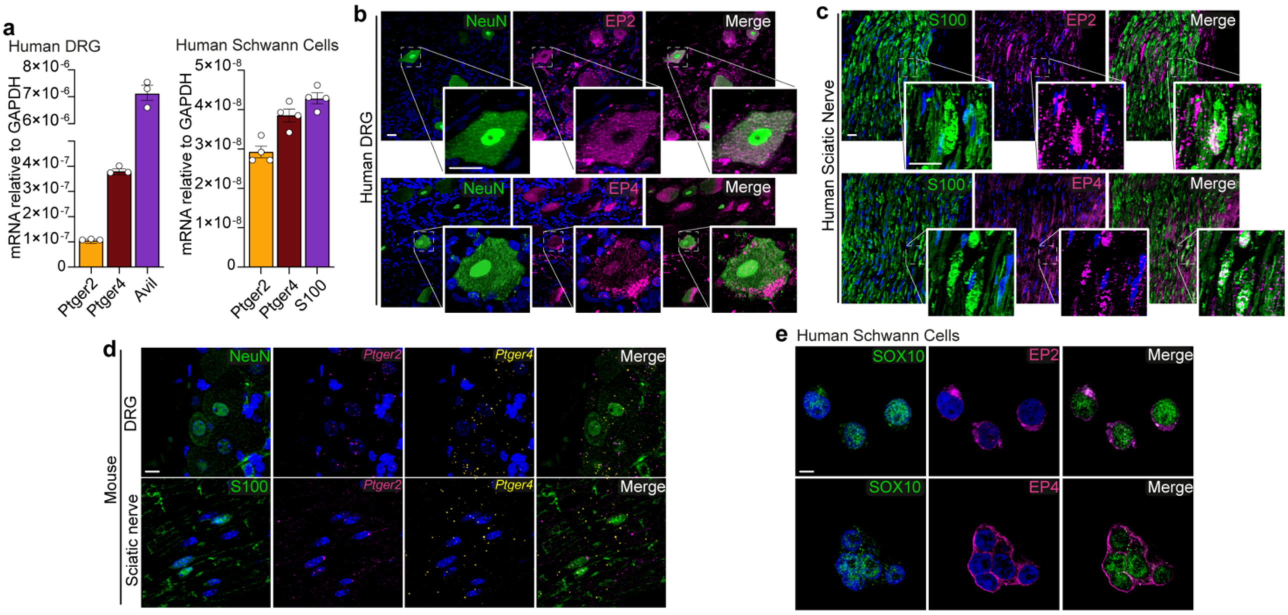

**Supplementary Fig. 2**. **EP2 and EP4 expression in human dorsal root ganglia and Schwann cells. a**, RT-qPCR for *Ptger2, Ptger4, Avil,* and *S100* mRNA in human dorsal root ganglia (DRG) and Schwann cells (SCs) (n=3 and 4 independent experiments, respectively). **b**, Representative images of EP2, EP4, NeuN and S100B expression in human DRG and **c**, sciatic nerve tissue (scale bar: 20 µm, inset 20µm) (n=3 subjects). **d**, Representative RNAscope images of *Ptger2* and *Ptger4* mRNA and S100 and NeuN protein expression in mouse DRG and sciatic nerve tissues (scale bar: 20 µm) (n=3 independent experiments). **e**, Representative images of EP2, EP4 and SOX10 expression in human SCs (n=3 independent experiments). Data are mean ± s.e.m.

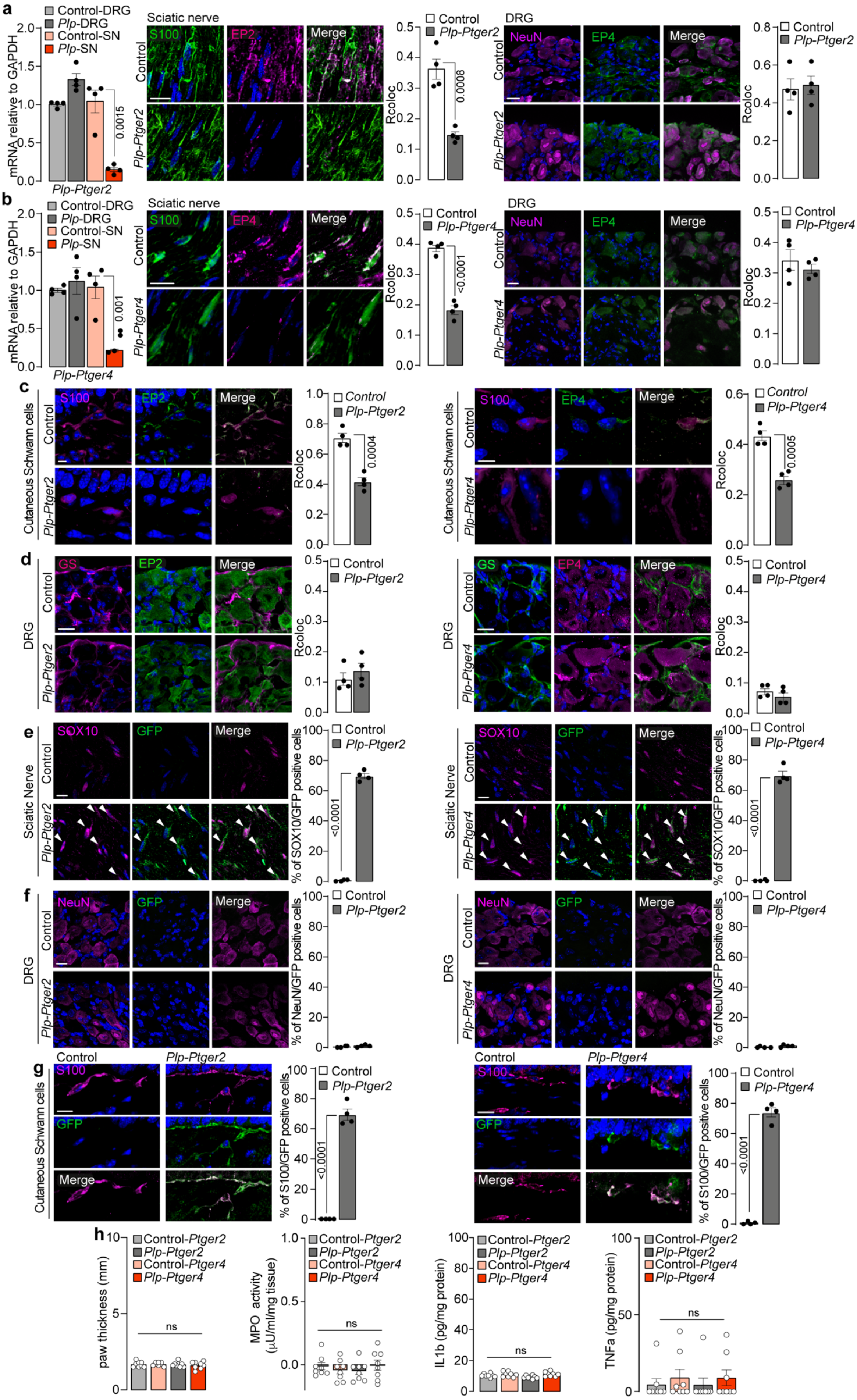
a
Control-DRG
Plp-DRG
Control-SN
Plp-SN
mRNA relative to GAPDH
Plp-Ptger2
0.0015
Sciatic nerve
S100
EP2
Merge
Control
Plp-Ptger2
Rcoloc
0.0008
DRG
NeuN
EP4
b
Plp-Ptger4
0.001
<0.0001
c
Cutaneous Schwann cells
0.0004
0.0005
d
GS
e
Sciatic Nerve
SOX10
GFP
% of SOX10/GFP positive cells
f
% of NeuN/GFP positive cells
g
% of S100/GFP positive cells
h
Control-Ptger2
Plp-Ptger2
Control-Ptger4
Plp-Ptger4
paw thickness (mm)
ns
MPO activity (μU/ml/mg tissue)
IL1b (pg/mg protein)
TNFa (pg/mg protein)

**Supplementary Fig. 3**. **AAV mediated EP2 and EP4 selective silencing in Schwann cells. a, b** RT-qPCR for *Ptger2* and *Ptger4* mRNA and representative images of EP2, EP4, NeuN and S100B protein expression and colocalization data (Rcoloc) in mouse sciatic nerve and dorsal root ganglia (DRG) in *P1p-Cre* and Control mice infected with AAV for selective silencing of EP2 (*Plp-Ptger2*) or EP4 (*Plp-Ptger4*). **c,** Representative images of EP2, EP4 and S100B protein expression and Rcoloc in mouse paw skin tissue of *P1p-Ptger2, P1p-Ptger4* and Control mice. **d,** Representative images of EP2, EP4 and satellite glial cell marker (GS) protein expression and Rcoloc in DRG of *P1p-Ptger2, P1p-Ptger4* and Control mice. **e, f, g,** Representative images of SOX10, NeuN, S100B and GFP protein expression and percentage of GFP positive cells in mouse sciatic nerve, DRG and paw skin tissue of *P1p-Ptger2*, *P1p-Ptger4* and Control mice (n=4 subjects). (scale bar: 20 μm) **h,** Paw thickness, myeloperoxidase (MPO) activity, IL-1β and TNF-α levels in paw tissue homogenates of *P1p-Ptger2, P1p-Ptger4* and Control mice. (n=8 mice per group). (ns, not significant). Data are mean ± s.e.m. **a, b, h,** 1-way or **a-g,** Student's t test.

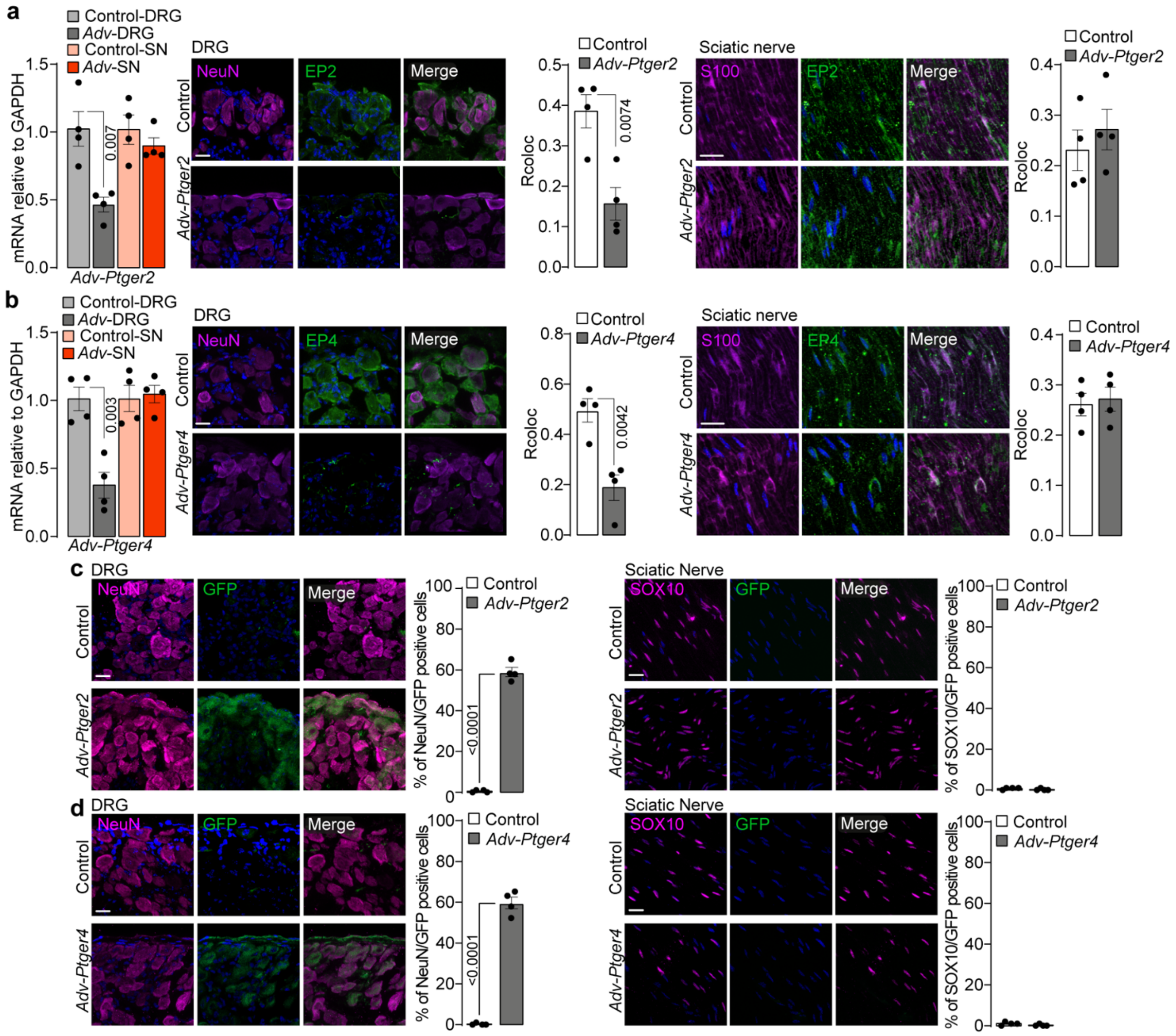

**Supplementary Fig. 4. AAV mediated EP2 and EP4 selective silencing in primary sensory neurons. a, b,** RT-qPCR for *Ptger2* and *Ptger4* mRNA and EP2, EP4, NeuN and S100B protein expression and colocalization data (Rcoloc) in mouse dorsal root ganglia (DRG) and sciatic nerve tissue in *Adv-Cre* and Control mice infected with AAV for selective silencing of EP2 (*Adv-Ptger2*) or EP4 (*Adv-Ptger4*). **c, d,** Representative images of NeuN, SOX10 and GFP protein expression and percentage of GFP positive cells in mouse DRG and sciatic nerve tissue of *Adv-Ptger2*, *Adv-Ptger4* and Control mice. (scale bar: 20 µm) (n=4 subjects). Data are mean ± s.e.m. **a, b,** 1-way or **a-d,** Student's t test.

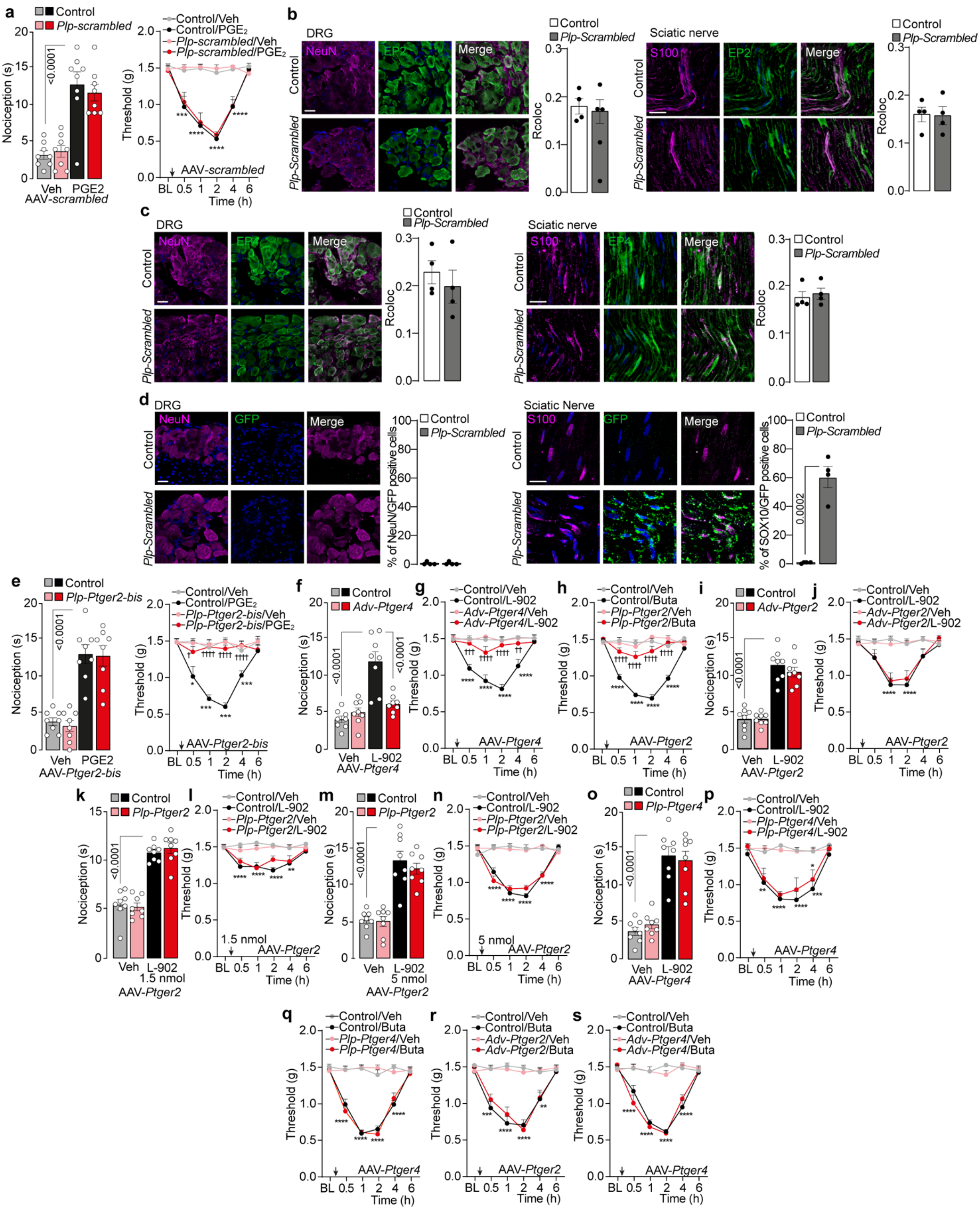

**Supplementary Fig. 5**. **Acute nociception and mechanical allodynia (allodynia) mediated by EP2 and EP4 selective agonist in mice with EP2 and EP4 selective deletion**. **a,** Acute nociception and allodynia after intraplantar (i.pl.) injection of $PGE_2$ (1.5 nmol) or vehicle (Veh) in *P1p-Cre* and Control mice infected with AAV for a scrambled shRNA (*Plp-*

*scrambled*). (n=8 mice per group). **b, c,** Representative images of EP2, EP4, NeuN and S100B protein expression and colocalization data (Rcoloc) in mouse sciatic nerve and dorsal root ganglia (DRG) of *P1p-scrambled* and Control mice. (scale bar: 20 μm) **d,** Representative images of NeuN, S100B and GFP protein expression and percentage of GFP positive cells in mouse DRG and sciatic nerve tissue of *P1p-scrambled* and Control mice. (scale bar: 20 μm) **e,** Acute nociception, and allodynia after i.pl. $PGE_2$ (1.5 nmol) or Veh in *P1p-Cre* and Control mice infected with AAV for a selective silencing of EP2 (*Plp-Ptger2-bis*). **f, i, k, m, o,** Acute nociception, and **g, j, l, n, p,** allodynia after i.pl. L-902,688 (L-902, 5 nmol) or Veh in **f, g,** *Adv-Ptger4* **i, j,** *Adv-Ptger2,* **k, l, m, n,** *Plp-Ptger2,* **o, p,** *Plp-Ptger4* and Control mice. (n=4 subjects) Allodynia after i.pl. Butaprost (Buta, 1.5 nmol) or Veh in **h,** *Plp-Ptger2,* **q,** *Plp-Ptger4,* **r,** *Adv-Ptger2,* **s,** *Adv-Ptger4* and Control mice. (n=8 mice per group). Data are mean ± s.e.m. **a, e, f, i, k, m, o**, 1-way or **a, e, g, h, j, l, n, p-s** 2-way ANOVA, Bonferroni correction, **b,c, d** Student's t test. *P<0.05, **P<0.01, ***P<0.001, ****P<0.0001 vs. Veh $^{†}$P<0.05, $^{††}$P<0.01, $^{†††}$P<0.001, $^{††††}$P<0.0001 vs. Buta, L-902, $PGE_2$/Veh.

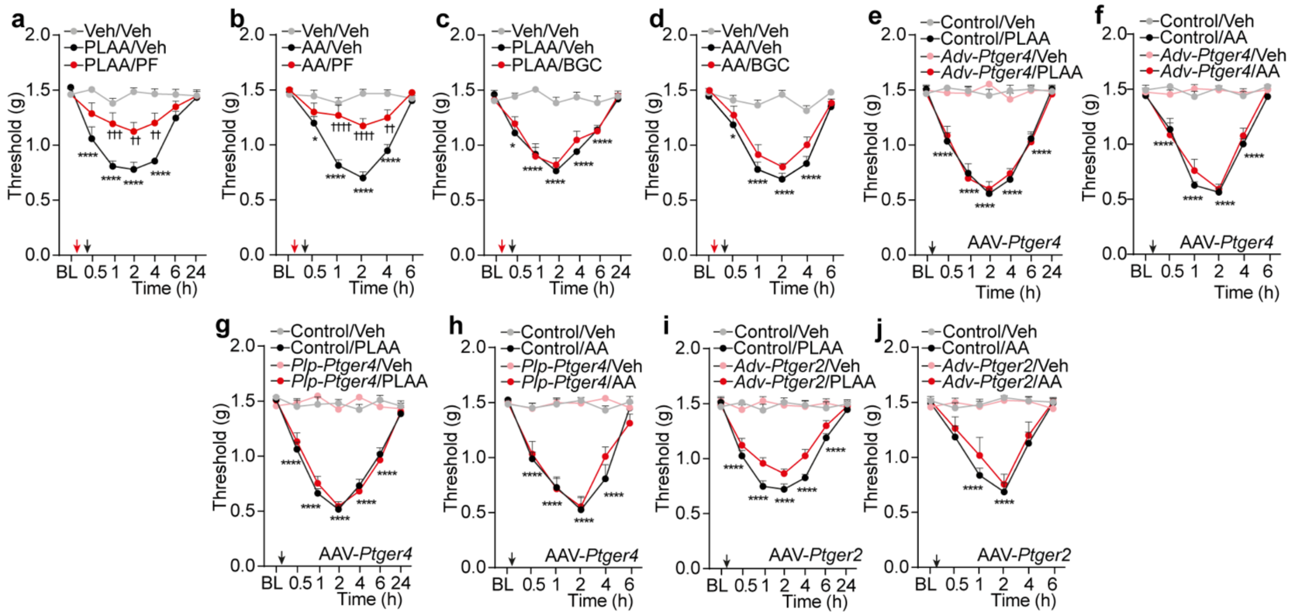

**Supplementary Fig. 6. EP2 in Schwann cells mediates hind paw mechanical allodynia (allodynia) by Phospholipase A2 Activating Protein (PLAA) and Arachidonic Acid (AA). a-d,** Allodynia after intraplantar (i.pl.) PLAA (2 nmol), AA (10 nmol) or vehicle (Veh) in C57BL/6J (B6) mice pretreated with PF-04448948 (PF, 5 nmol), BGC-20-1531 (BGC, 5 nmol) or Veh (n=8 mice per group). **e-j,** Allodynia after i.pl. PLAA, AA or Veh in *P1p-Cre*, *Adv-Cre* or Control mice infected with AAV for selective silencing of EP2 (-Ptger2) or EP4 (-Ptger4) (*Plp-Ptger2* or *Adv-Ptger2* and *Plp-Ptger4* or *Adv-Ptger4*) (n=8 mice per group). Data are mean ± s.e.m. 2-way ANOVA, Bonferroni correction. *P<0.05, ****P<0.0001 vs. Veh, Control/Veh, $^{\dagger\dagger}$P<0.01, $^{\dagger\dagger\dagger}$P<0.001, $^{\dagger\dagger\dagger\dagger}$P<0.0001 vs. PLAA, AA/Veh.

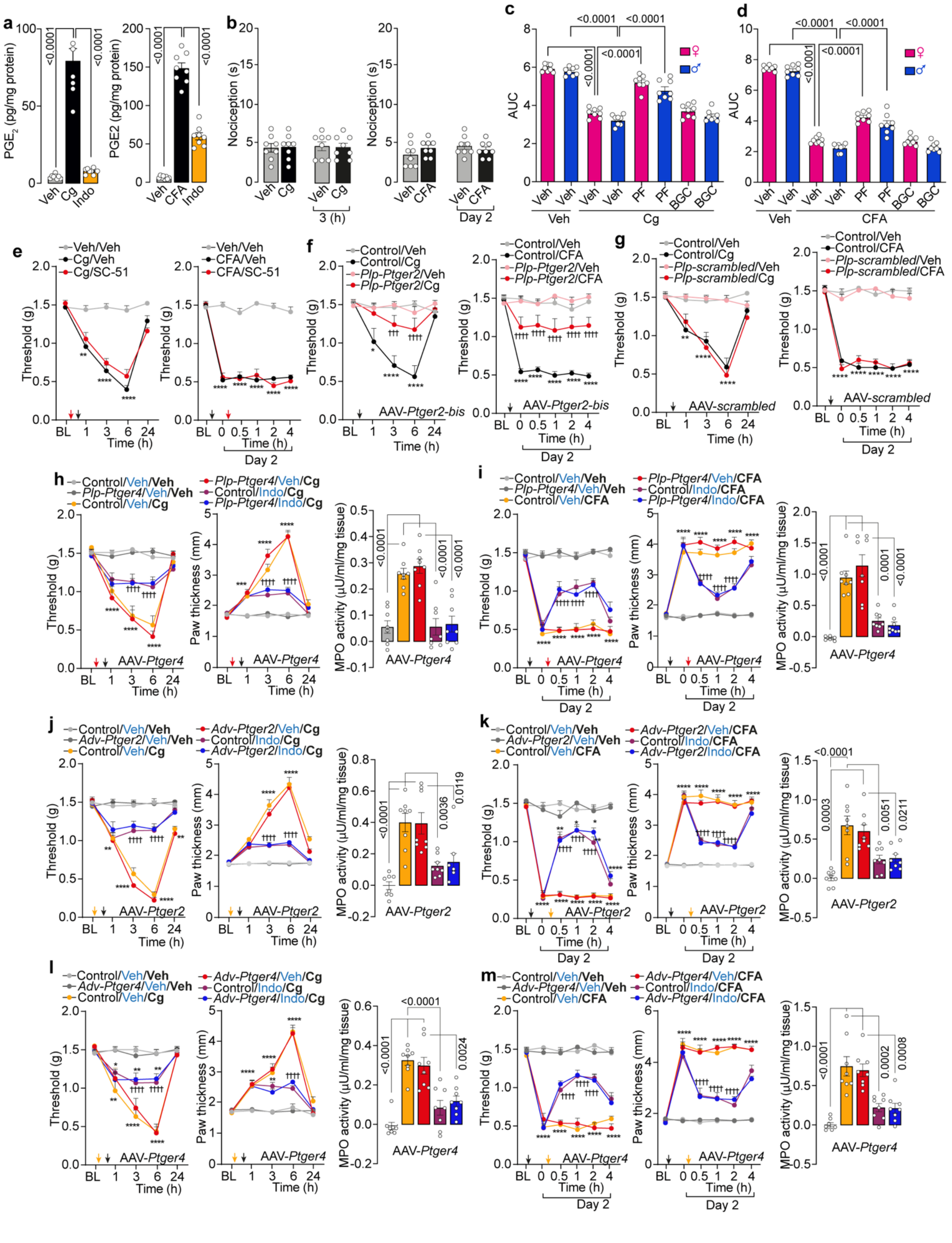

**Supplementary Fig. 7. EP2 in Schwann cells mediates carrageenan and CFA hind paw mechanical allodynia (allodynia) but not inflammation. a,** $PGE_2$ assay in paw tissue homogenates of C57BL/6J (B6) mice after intraplantar (i.pl.) carrageenan (Cg, 300 µg) or Complete Freud Adjuvant (CFA) or vehicle (Veh) pretreated with indomethacin

(Indo, 280 nmol) or Veh. **b,** Acute or delayed nociception after i.pl. Cg, CFA or Veh in B6 mice. **c,d** Allodynia expressed as area under the curve (AUC) after i.pl. Cg (300 μg), CFA or Veh in male and female B6 mice pretreated with PF-04448948 (PF, 5 nmol), BGC-20-1531 (BGC, 5 nmol) or Veh. **e,** Allodynia after i.pl. Cg, CFA or Veh in B6 mice pretreated with SC-51322 (SC-51, 5 nmol) or Veh. **f,g,** Allodynia after i.pl. Cg, CFA or Veh in *P1p-Cre* or Control mice infected with AAV for selective silencing of EP2 (*Ptger2 bis*) or scrambled (*P1p-Ptger2-bis* or *P1p-scrambled*). **h-m,** Allodynia, paw thickness and MPO activity assay after i.pl. Cg, CFA or Veh in *P1p-Cre*, *Adv-Cre* or Control mice infected with AAV for selective silencing of EP2 (-Ptger2) or EP4 (-Ptger4) (*P1p-Ptger2* or *Adv-Ptger2* and *P1p-Ptger4* or *Adv-Ptger4*) and pretreated with indomethacin (Indo, 280 nmol) or Veh (n=8 mice per group). Data are mean ± s.e.m. **a, c, d h-m** MPO activity 1-way or **e-m** allodynia and paw thickness 2-way ANOVA, Bonferroni correction. *P<0.05, **P<0.01 ***P<0.0001, ****P<0.0001 vs. Control/Veh/Veh or Veh/Veh; $^{††††}$P<0.0001vs. Control/Veh/Cg, CFA or Cg, CFA/Veh.

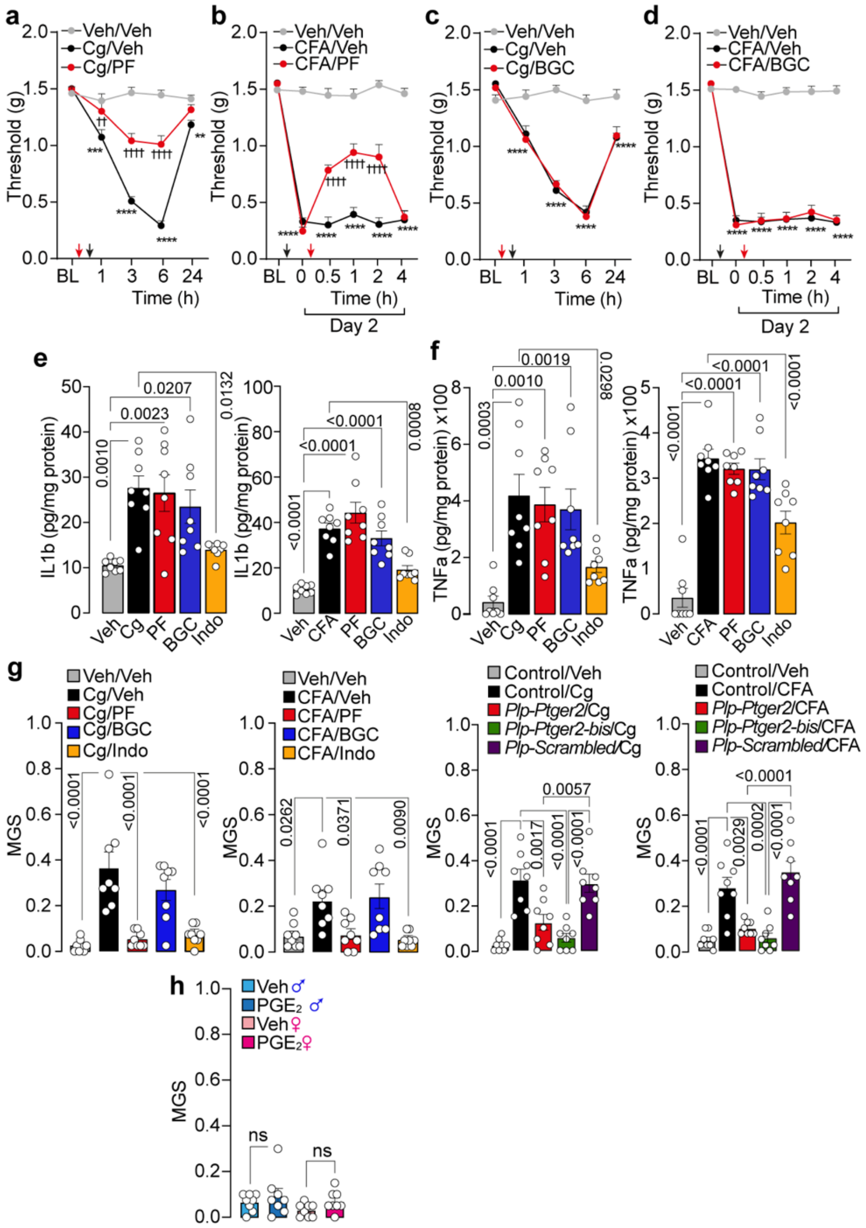

**Supplementary Fig. 8 EP2 in Schwann cells mediates carrageenan and CFA hind paw mechanical allodynia (allodynia) and grimace behaviour but not inflammation. a-d** Allodynia after intraplantar (i.pl.) carrageenan (Cg, 300 µg) or Complete Freud Adjuvant (CFA) or vehicle (Veh) in C57BL/6J (B6) mice pretreated with with PF-04448948 (PF, 5 nmol), BGC-20-1531 (BGC, 5 nmol) or Veh. (n=8 mice per group). **e, f,** IL1β and TNFα assay in paw tissue homogenates after i.pl. Cg, CFA or Veh in B6 mice pretreated with PF (5 nmol), BGC (5 nmol) indomethacin (Indo, 280 nmol) or Veh. **g,** mouse grimace score (MGS) after i.pl. Cg, CFA or Veh in B6 mice pretreated with PF (5 nmol), BGC (5 nmol), Indo (280 nmol) or Veh or in *P1p-Cre* or Control mice infected with AAV for selective silencing of EP2 (*Ptger2 and Ptger bis*) or scrambled (*P1p-Ptger2, P1p-Ptger2-bis* or *P1p-scrambled*). **h,** MGS after i.pl. $PGE_2$ in B6 male and female mice (n=8 mice per group). Data are mean ± s.e.m. **a-d** 2-way or **e-g** 1-way ANOVA, Bonferroni correction. **P<0.01 ***P<0.0001, ****P<0.0001 vs. Veh/Veh; $^{\dagger\dagger}$P<0.01, $^{\dagger\dagger\dagger\dagger}$P<0.0001vs. Cg/Veh, CFA/Veh.

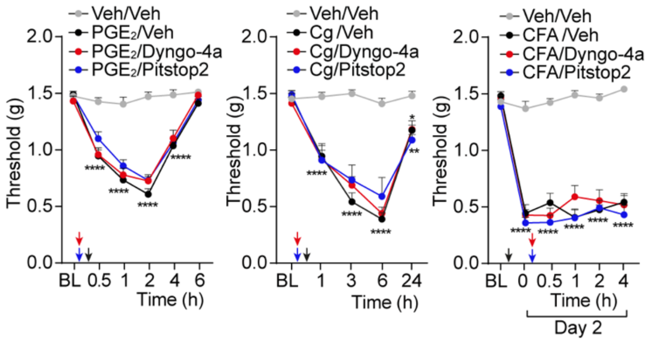

**Supplementary Fig. 9. Clathrin and dynamin inhibitors do not mediate $PGE_2$, carrageenan and CFA hind paw mechanical allodynia (allodynia).** Allodynia after intraplantar (i.pl.) injection of $PGE_2$ (1.5 nmol), carrageenan (Cg, 300 µg), Complete Freud Adjuvant (CFA) or vehicle (Veh) in C57BL/6J (B6) mice pretreated with Dyngo-4a (500 pmol), Pitstop2 (500 pmol) or Veh (n=8 mice per group). Data are mean ± s.e.m. 2-way ANOVA, Bonferroni correction. *P<0.05, **P<0.01, ****P<0.0001 vs. Veh/Veh.

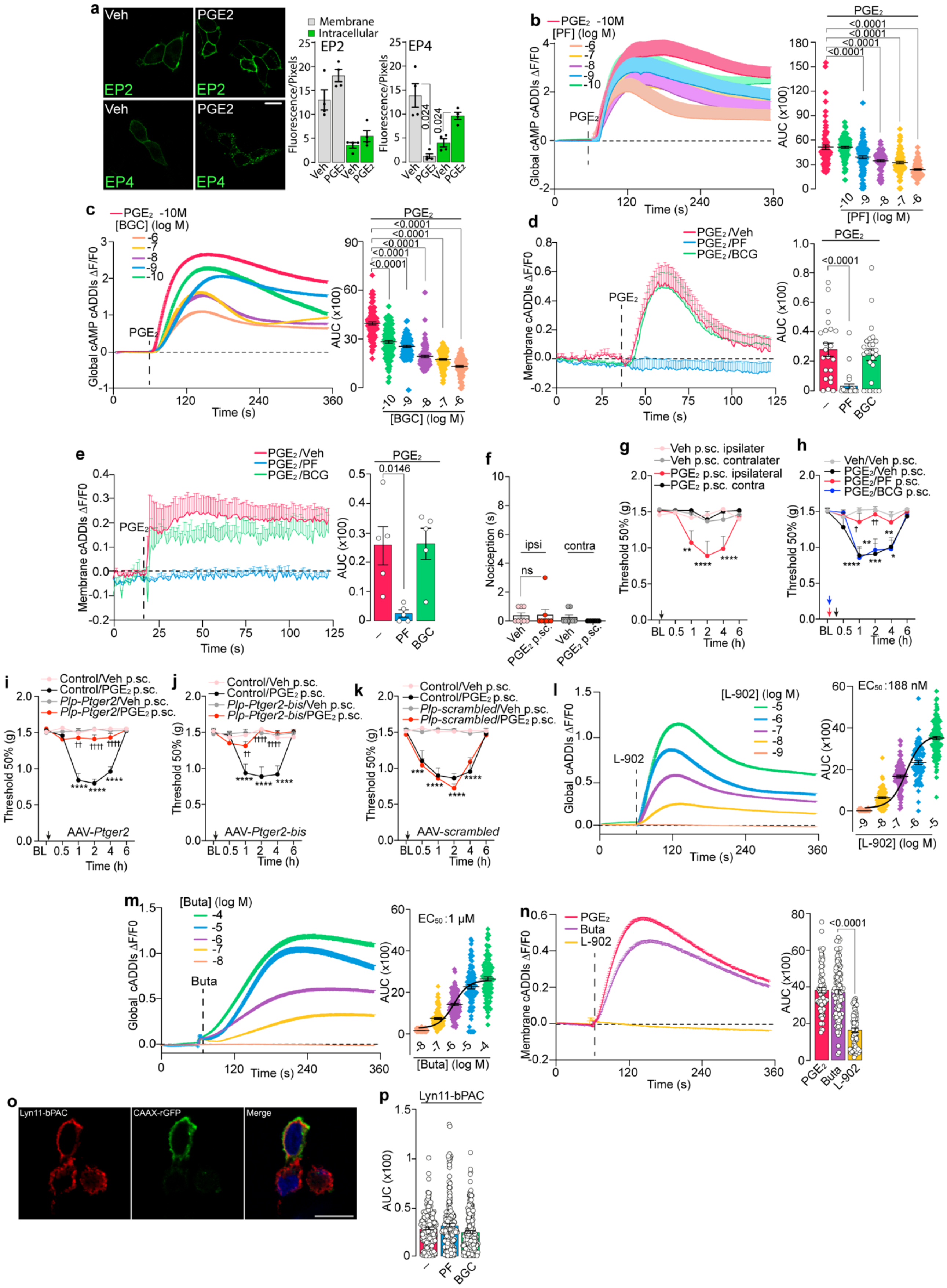

**Supplementary Fig. 10. Schwann cells cAMP nanodomains promote EP2 hind paw mechanical allodynia (allodynia). a,** Representative images and cumulative data of membrane and intracellular localization of EP2 and EP4 in HEK293T cells, with or without pre-incubation with unlabeled PGE2 (10 µM, 30 min) (n=4 independent experiments). (Scale bar, 20 µm). **b,c,** Global cAMP measurement after $PGE_2$ (100 pM) stimulation of human Schwann cells (hSCs) in the presence of PF-04448948 (PF) (cells number: -6 M=111, -7M=89, -8 M=75, -9M=99, -10M=99), BGC-20-1531 (BGC) (cells number: -6 M=96, -7M=102, -8 M=87, -9M=84, -10M=95) or Veh (cells number: n=76 or n=91) (n=3 independent experiments). **d,e,** Membrane cAMP measurement after $PGE_2$ (10 nM) stimulation of mouse sciatic nerve and cutaneous SCs in the presence of PF (1 µM) (cells number: sciatic nerve SCs n=27, cutaneous SCs n=5), BGC (1 µM) (cells number: sciatic nerve SCs n=26, cutaneous SCs n=5 ) or Veh (cells number: sciatic nerve SCs n=22, cutaneous SCs n=5) (n=3 independent experiments). **f-h** Acute nociception and allodynia after peri-sciatic (p.sc.) injection of $PGE_2$ (1.5 nmol) or vehicle (Veh) in C57BL/6J mice (B6) pretreated with PF-04448948 (PF, 5 nmol), BGC-20-1531 (BGC, 5 nmol) or Veh. **i,j,k,** Allodynia after p.sc. $PGE_2$ (1.5 nmol) or vehicle (Veh) in *P1p-Cre* or Control mice infected with AAV for selective silencing of EP2 (*Ptger2 and Ptger bis*) or scrambled (*P1p-Ptger2, P1p-Ptger2-bis* or *P1p-scrambled*). (n=8 mice per group). **l,m,** Global cAMP measurement after L-902,688 (L-902) (cells number: -5M=112,-6M=73,-7M=94,-8M=82,-9M=94, n=3 independent experiments) and (R)-Butaprost (Buta) (cells number: -4M=109,-5M=96,-6M=99,-7M=94,-8M=98, n=3 independent experiments) stimulation of hSCs. **n,** Membrane cAMP measurement after $PGE_2$ (10 nM), Buta (100 µM) or L-902 (100 µM) stimulation of hSCs (cells number: $PGE_2$=83, Buta=104, L-902=79, n=3 independent experiments). **o**, Colocalization of Lyn11-bPAC and the plasma membrane marker CAAX in HEK293 cells (scale bar: 10 µm). **p,** Membrane cAMP formation in hSCs after blue light (450 nm, 10 min) activation of membrane-tagged bPAC (Lyn11-bPAC) in the presence of PF (1 µM) (cells number: n=214,), BGC (1 µM) (cells number: n=214) or Veh (cells number: n=214,). Data are mean ± s.e.m. **a** Student's t test**, b-f, n,p** 1-way, **j-k** 2-way ANOVA Bonferroni correction. AUC, area under the curve. *P<0.05, **P<0.01 ***P<0.0001, ****P<0.0001 vs. Veh, Control/Veh, $^{†}$P<0.05, $^{††}$P<0.01, $^{††††}$P<0.0001 vs. Control/$PGE_2$.

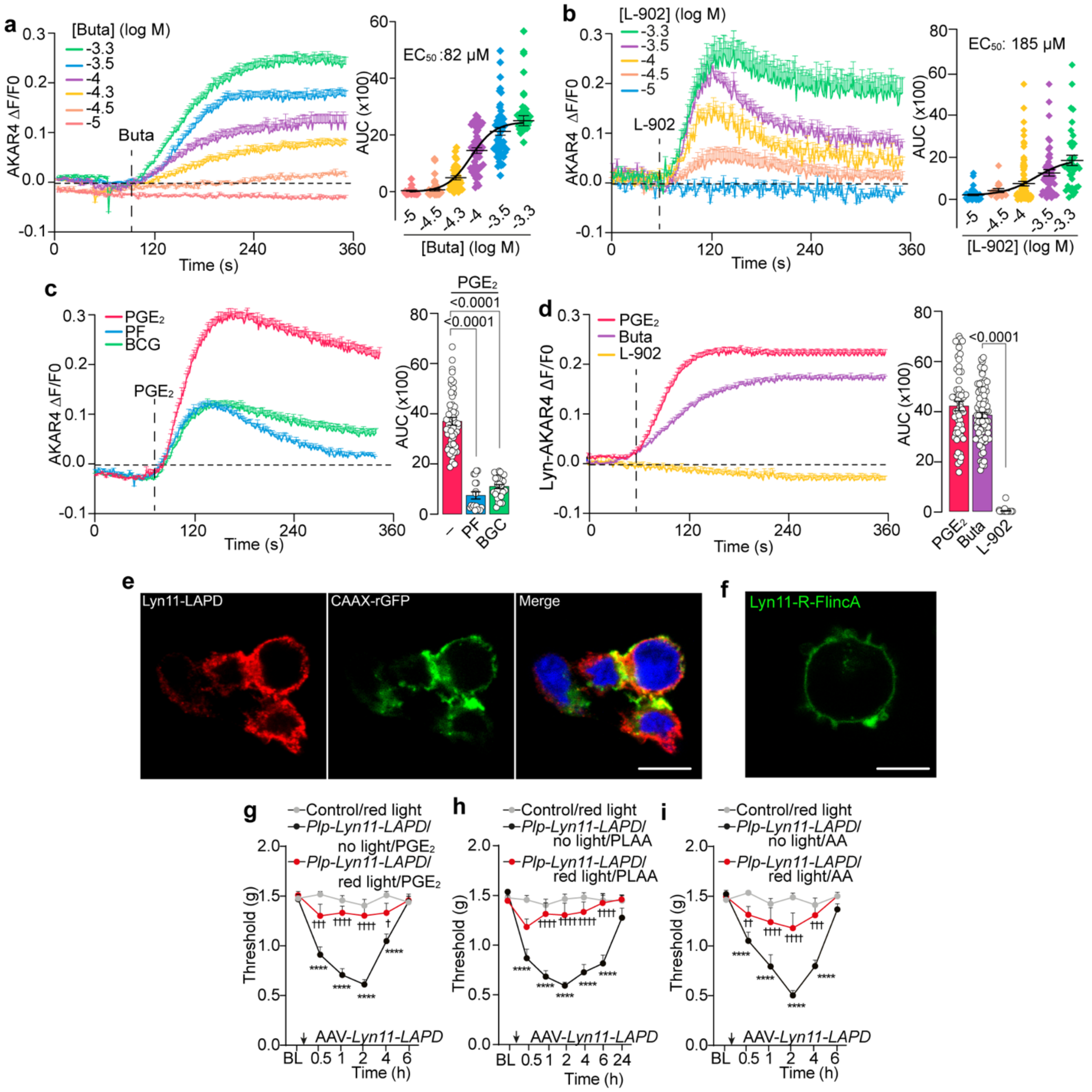

**Supplementary Fig. 11. Schwann cells membrane-bound PKA/AKAP drives EP2 signaling. a,b** Global PKA activation induced by Buta (cells number:-3.3M=37,-3.5M=48,-4M=49,-4.3M=35,4.5M=44,-5M=37, n=3 independent experiments) and L-902 (cells number:-3.3M=37,-3.5M=52,-4M=95,-4.5M=16,-5M=51, n=3 independent experiments) stimulation of human Schwann cells (hSCs). **c**, global PKA activation induced by $PGE_2$ (10 nM) in the presence of PF (1 μM), BGC (1 μM) or Veh (cells number: PF=18, BGC=29, $PGE_2$=62, n=3 independent experiments) in hSCs. **d**, Membrane PKA activation induced by $PGE_2$, (10 nM) Buta (100μM) or L-902 (100 μM) stimulation of hSCs (cells number: $PGE_2$=57, Buta=81, L-902=45, n=3 independent experiments). **e**, Colocalization of Lyn11-LAPD and the plasma membrane marker CAAX in HEK293 cells. (scale bar: 10 μm). **f,** Membrane localization of Lyn11-R-FlincA in HEK293 cells (scale bar: 10 μm). **g,h,i,** Allodynia after i.pl. $PGE_2$,

Phospholipase A2 Activating Protein (PLAA, 2 nmol), Arachidonic Acid (AA, 10 nmol) or Veh in Plp-Cre or Control mice after i.pl. infection with AAV-Lyn11-LAPD (Plp- Lyn11-LAPD) and stimulated with red light (n=8 mice per group). Data are mean ± s.e.m. **c,d,** 1-way or**, g-i,** 2-way ANOVA, Bonferroni correction. AUC, area under the curve. ***P<0.0001 vs. Veh, Control/red light. †P<0.05, ††P<0.01, †††P<0.001, ††††P<0.0001 vs. Plp-Lyn11-LAPD/no light/$PGE_2$.

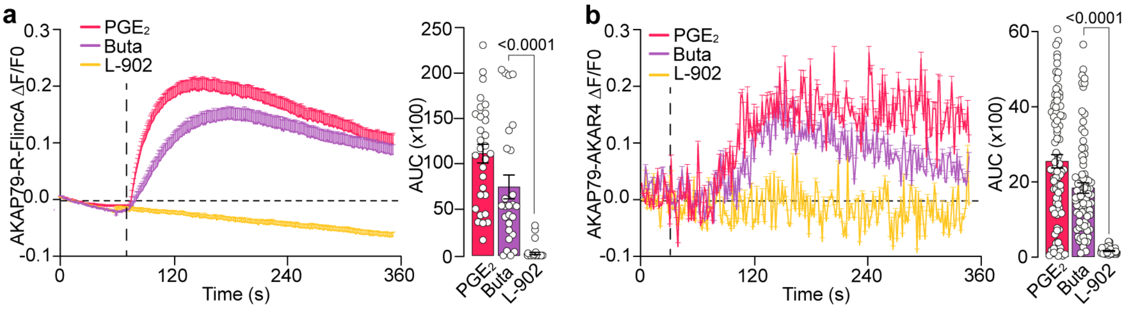

**Supplementary Fig. 12. Schwann cells membrane-bound PKA/AKAP drives EP2 signaling. a,** AKAP79 associated cAMP formation after $PGE_2$ (10 nM), (R)-Butaprost (Buta, 100µM) or L-902,688 (L-902, 100 µM) stimulation of human Schwann cells (hSCs) (cells number: $PGE_2$=30, Buta=26, L-902=37, n=3 independent experiments). **b,** AKAP79 associated PKA activation induce by $PGE_2$ (10 nM), Buta (100 µM) or L-902 (100 µM) stimulation of hSCs (cells number: $PGE_2$=86, Buta=82, L-902=69, n=3 independent experiments). AUC, area under the curve. Data are mean ± s.e.m. 1-way ANOVA, Bonferroni correction.

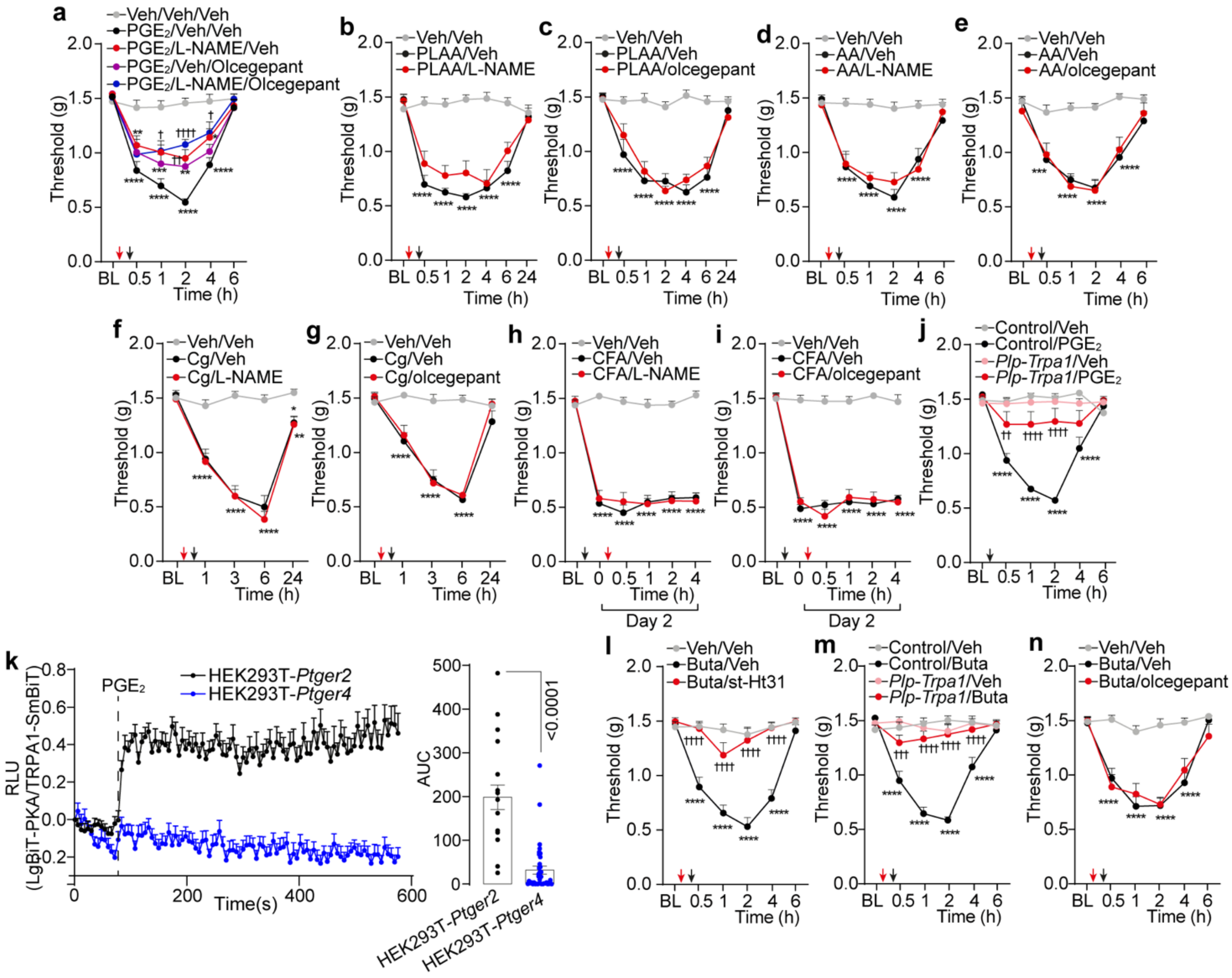

**Supplementary Fig. 13. EP2 associates with TRPA1 in Schwann cells signals hind paw mechanical allodynia (allodynia). a,** Allodynia after intraplantar (i.pl.) injection of $PGE_2$ (1.5 nmol) or vehicle (Veh) in C57BL/6J (B6) mice pretreated with L-NAME (1 µmol), olcegepant (1 nmol), combination of L-NAME + olcegepant or Veh (n=8 mice per group). **b-i,** Allodynia after i.pl. Phospholipase A2 Activating Protein (PLAA, 2 nmol), Arachidonic Acid (AA, 10 nmol), carrageenan (Cg, 300 µg), Complete Freud Adjuvant (CFA) or Veh in B6 mice pretreated with L-NAME (1 µmol), olcegepant (1 nmol) or Veh (n=8 mice per group). **j,** Allodynia after i.pl. of $PGE_2$ or Veh in *Plp-Trpa1* or Control mice (n =8 mice per group). **k,** Effect of $PGE_2$ (100 nM) stimulation in catalytic subunit of PKA and TRPA1 interaction in HEK293T cells transfected with the EP2 (-Ptger2) or EP4 (-Ptger4). **l,** Allodynia after i.pl. butaprost (Buta, 1.5 nmol) or Veh in B6 mice pretreated with st-Ht31 (17 nmol) or Veh (n=8 mice per group). **m,** Allodynia after i.pl. Buta in *Plp-Trpa1* or Control mice. **n,** Allodynia after i.pl. Buta in B6 mice pretreated with olcegepant (1 nmol) (n =8 mice per group). Data are mean ± s.e.m. **k,** 1-way or **a-j, l-n,** 2-way ANOVA, Bonferroni correction. AUC, area under the curve.

*P<0.05, **P<0.01, ***P<0.001, ****P<0.0001 vs. Veh, Control/Veh $^{\dagger}$P<0.05, $^{\dagger\dagger}$P<0.01, $^{\dagger\dagger\dagger}$P<0.001, $^{\dagger\dagger\dagger\dagger}$P<0.0001 vs. $PGE_2$, PLAA, AA, Cg, CFA, Buta/Veh, or Control/$PGE_2$, Buta.

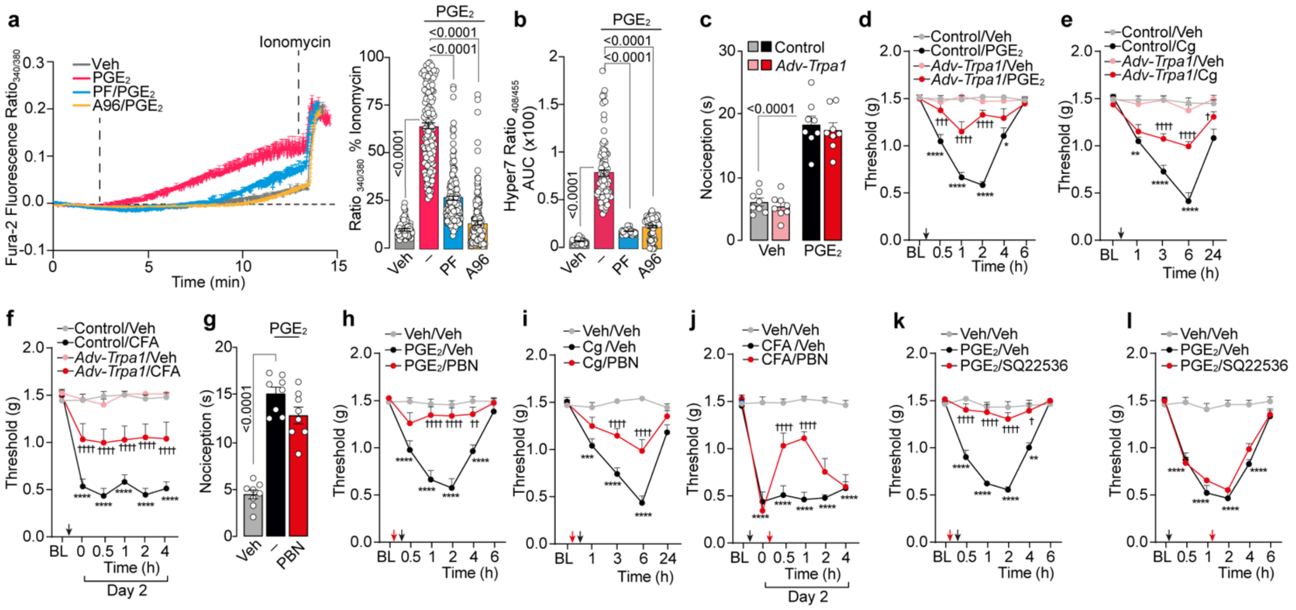

**Supplementary Fig. 14. TRPA1 in Schwann cells sustains $PGE_2$ hind paw mechanical allodynia (allodynia).** **a,** Typical traces and cumulative data of calcium response after $PGE_2$ stimulation of human Schwann cells (hSCs) and in the presence of PF-04448948 (PF), A967079 (A96) or vehicle (Veh) (cells number: $PGE_2$=174, PF=156, A96=224, Veh=99, n=3 independent experiments). **b,** Cumulative data of HyPer7.2, after $PGE_2$ stimulation of hSCs, PF, A96 or Veh (cells number: $PGE_2$=86, PF=63, A96=89, Veh= 63, n=3 independent experiments). **c, d,** Acute nociception and allodynia after intraplantar (i.pl.) injection of $PGE_2$ (1.5 nmol) or Veh in *Adv-Trpa1* or Control mice. **e, f** allodynia after carrageenan (Cg, 300 µg), Complete Freud Adjuvant (CFA) or vehicle (Veh) in *Adv-Trpa1* or Control mice. **g,h** Acute nociception and allodynia after i.pl. $PGE_2$ or Veh in C57BL/6J (B6) mice pretreated with PBN (1 µmol); **i,j,** allodynia after i.pl. Cg, CFA or Veh in B6 mice treated with PBN (1 µmol). **k,l,** allodynia after i.pl $PGE_2$ or Veh B6 mice pre or post treated with SQ25536 (25 nmol). (n =8 mice per group). Data are mean ± s.e.m. **a,b,c,g** 1-way or **d-f, h-l,** 2-way ANOVA, Bonferroni correction. AUC, area under the curve. *P<0.05, **P<0.01, ****P<0.0001 vs. Veh, Control/Veh †P<0.05, ††P<0.01, †††P<0.001, ††††P<0.0001 vs. Control $PGE_2$, Control Cg, Control CFA, $PGE_2$, Cg, CFA.

**Supplementary Table 1. Primers for plasmid construction.**

| Primer | Sequence (5' to 3') |
|---|---|
| P1 | AAAGCTAGCATGTTCACCTTGGGATAC |
| P2 | AAAGAGCTCTCTGAAACTCAGAAAACTCC |
| P3 | AAAGAGCTCAAATGTTCACCTTGGGATAC |
| P4 | AAATCTAGACTAAAACTCAGAAAACTCC |
| P5 | AAAGCGATCGCATGAAGCGCAGCCTGAG |
| P6 | AAACTCGAGTTCTGAGGCTCAAGATGGTGTGT |
| P7 | AAACTCGAGAATGAAGCGCAGCCTGAG |
| P8 | AATCTAGACTAAGGCTCAAGATGGTGTGT |
| P9 | CTGCAGCGACCCGCTTAAAAATGTTCACCTTGGGATACCATG |
| P10 | CCGCTCGAGCCAGAATTCCCCTGAAACTCAGAAAACTCCTTGC CAC |
| P11 | TGGTATCCCAAGGTGAACATTTTTAAGCGGGTCGCTGCAG |
| P12 | AGGAGTTTTCTGAGTTTCAGGGGAATTCTGGCTCGAG |
| P13 | TGTTCGAGGAGATTCTCGGGATGAAGCGCAGCCTGAGG |
| P14 | TAGAAGATCTGCTAGCTTAGCTAAGGCTCAAGATGGTGTGTTTT TG |
| P15 | TTCCTCAGGCTGCGCTTCATCCCGAGAATCTCCTCGAACAG |
| P16 | CACACCATCTTGAGCCTTAGCTAAGCTAGCAGATCTTCTAGAGT CGG |
| P17 | GAGTAACCATCAACAGTGGGATGTTCACCTTGGGATACCATGG |
| P18 | GGCCGCCCCGACTCTAGAAGCTAAAACTCAGAAAACTCCTTGC CACAC |
| P19 | TGGTATCCCAAGGTGAACATCCCACTGTTGATGGTTACTCG |
| P20 | AGGAGTTTTCTGAGTTTTAGCTTCTAGAGTCGGGGCGGCC |
| P21 | TGGGAGCTCAGGGGAATTCTATGAAGCGCAGCCTGAGGA |
| P22 | AGCCGGTAGCCGGTCACACCATAAGGCTCAAGATGGTGTGTTT TTGC |
| P23 | TTCCTCAGGCTGCGCTTCATAGAATTCCCCTGAGCTCCCAC |
| P24 | CACACCATCTTGAGCCTTATGGTGTGACCGGCTACCG |
| P25 | AAAGCTAGCGGATCCATGATGAAGCGGCTGGT |
| P26 | AAGAATTCGCGATCGCCTACTTGTCGTTTTCCAGGGT |
| P27 | CTAGCGCCACCATGGGATGTATCAAATCGAAAGGTAAG |
| P28 | GACAGCCTGTCAGGAAGTGGTTGCATTAAGAGCAAGGGAAAG GATTCGTTCTGCTATGAAAATGAAGTTGCCG |
| P29 | GATCCGGCAACTTCATTTTCATAGCAGAACGAATCCTTTCC |
| P30 | TTGCTCTTAATGCAACCACTTCCTGACAGGCTGTCCTTACCTTT CGATTTGATACATCCCATGGTGGCG |
| P31 | CACGCTAGCATGGAAACCACAATTTCAG |
| P32 | GGGGGATCCCTGTAGAAGATTGTTTATTTT |
| P33 | CTAGGATGGGATGTATCAAATCGAAAGGT |
| P34 | AAGGACAGCCTGTCAGGAAGTGGTTGCATTAAGAGCAAGGGA AAGGATTCGGTT |
| P35 | ACCACTTCCTGACAGGCTGTCCTTACCTTTCGATTTGATACATC CCATC |
| P36 | AACCGAATCCTTTCCCTTGCTCTTAATGCA |
| P37 | GACTGCTAGCATGAGCAGGGACCCCCTG |

| | |
|---|---|
| P38 | TCAGTCTAGATCAGTACGCTCAGCATCAAGGCTGC |
| P39 | ACTGGCTAGCATGGAAACCAC |
| P40 | GACTGCTAGCCTGTAGAAGATTG |
| P41 | CTAGCATGGGATGTATCAAATCGAAAGGTAAGGACAGCCTGTC |
| P42 | AGGAAGTGGTTGCATTAAGAGCAAGGGAAAGGATTCGTTCTGC TATGAAAATGAAGTTGCCG |
| P43 | CTAGCGGCAACTTCATTTTCATAGCAGAACGAATCCTTTCCCT |
| P44 | TGCTCTTAATGCAACCACTTCCTGACAGGCTGTCCTTACCTTTC GATTTGATACATCCCATG |
| P45 | CGCGCCACTAGTGTTAACCATATGCCTAGGTGCGATCGCTTAC GTATG |
| P46 | CTAGCATACGTAAGCGATCGCACCTAGGCATATGGTTAACACT AGTGG |
| P47 | AAACACGCGTTTACGGGGTCATTAGTTC |
| P48 | TCCGGTACCCATGGTGGCGGTTCACT |
| P49 | AAAGTTAACGCCACCATGATGAAGCGGCTGGT |
| P50 | AAAACTAGTTTACTTGTACAGCTCGTCCATGCCG |
| P51 | AAGTTAACATGAGCAGGGACCCCCTGCCCTT |
| P52 | AAACTAGTTTACTTGTCATCGTCGTCCTTGTAATCTGTACGCTC AGCATC |
| P53 | AAGCTAGCATGGATTACAAGGACGACGATGACAAGATGAGCA GGGACCCCCT |
| P54 | AAATACGTAGTACGCTCAGCATCAAGGCTGCAG |
| P55 | GGGCCTAGGCTGTAGAAGATTGTTTA |
| P56 | GGCACGCTTTCACTATGACCTTCTTTTACATCTGTGGCTTCACT AAAAGAAGGTCATAGTGAAAGCT |
| P57 | AGCGAGCTTTCACTATGACCTTCTTTTAGTGAAGCCACAGATGT AAAAGAAGGTCATAGTGAAAGCG |
| P58 | GGCAAGTGTTCATTAACCAGTTATATTACATCTGTGGCTTCACT AATATAACTGGTTAATGAACACG |
| P59 | AGCGCGTGTTCATTAACCAGTTATATTAGTGAAGCCACAGATG TAATATAACTGGTTAATGAACACT |
| P60 | GGCAGTCCTCATTCCGTTCGACTGTTTACATCTGTGGCTTCACT AAACAGTCGAACGGAATGAGGAT |
| P61 | AGCGATCCTCATTCCGTTCGACTGTTTAGTGAAGCCACAGATGT AAACAGTCGAACGGAATGAGGAC |
| P62 | GGCAGAATCCCAATGGTTACATCTAATACATCTGTGGCTTCACT ATTAGATGTAACCATTGGGATTT |
| P63 | AGCGAAATCCCAATGGTTACATCTAATAGTGAAGCCACAGATG TATTAGATGTAACCATTGGGATTC |

**Supplementary Table 2. Human and mouse primers for RT-qPCR.**

| Gene | Sequence (5' to 3') |
|---|---|
| Human GAPDH (NM_002046) | F: CTGTTCGACACTCAGCCGCATC<br>R: GCGCCCAATACGACCAAATCCG |
| Human AVIL (NM_006576) | F: CCTCCTCCCTAAACTCCAAT<br>R: AGTGTTCTCGCTGCCATC |
| Human S100 (NM_006271) | F: CTAGACGAGAATGGAGACGGG<br>R: AATGTGGCTGTCTGCTCAACT |
| Human $PGE_2$ (NM_000956) | F: CTCATTCTCCTGGCTATCATGAC<br>R: CCCATTTTTCCTTTCGGGAAG |
| Human PGE4 (NM_000958) | F: TCATCTTACTCATTGCCACCTC<br>R: TCACAGAAGCAATTCGGATGG |
| Mouse GAPDH (NM_008084) | F: AATGGTGAAGGTCGGTGTG<br>R: GTGGAGTCATACTGGAACATGTAG |
| Mouse AVIL (NM_009635) | F: CTTCCTCGCTCAACTCCAA<br>R: GCATTGCCATCACAGAGAAG |
| Mouse S100 (NM_011309) | F: GAAAGACCTGCTACAAACTGAAC<br>R: TCCAGTTCCTTCATTACCTTGTC |
| Mouse PGE2 (NM_008964) | F: GAGGTTTCATCCATGTAGGCA<br>R: AGAGGAGAGAGGACTTCGATG |
| Mouse PGE4 | F: CTGATGTCTTTCACCACGTTTG<br>R: CATCTTACTCATCGCCACCTC |
